# Defining the Antigenic Topology and Prospective Binding Breadth of Vaccination-induced SARS-CoV-2 Neutralizing Antibodies

**DOI:** 10.64898/2026.03.03.709398

**Authors:** Deepika Jaiswal, Clara G. Altomare, Daniel C. Adelsberg, Iden A. Sapse, Gagandeep Singh, Florian Krammer, Viviana Simon, Ali H. Ellebedy, Goran Bajic

## Abstract

Antibodies that neutralize SARS-CoV-2 primarily target the viral spike glycoprotein, yet the breadth of these responses is continually challenged by viral evolution. While extensive structural studies have defined epitopes across the spike protein, how antibodies elicited by the initial mRNA vaccination campaigns perform against subsequently emerging variants remains an important question. Here, we structurally and functionally characterize a panel of early plasmablast-derived human monoclonal antibodies isolated following primary mRNA vaccination, targeting both the receptor-binding domain (RBD) and the N-terminal domain (NTD) of spike. Using cryo–electron microscopy, variant-binding analyses, and viral-fusion inhibition assays, we observe that antibodies directed against immunodominant regions of the RBD and NTD are highly potent but more frequently impacted by variant-associated mutations. In contrast, antibodies engaging a conserved hydrophobic pocket within the NTD exhibit broader reactivity and neutralize through distinct molecular mechanisms. Together, these findings extend prior structural studies of spike-directed antibodies by prospectively assessing the breadth of vaccine-elicited antibodies against later variants and identifying structural features associated with differential escape sensitivity. These results contribute to a growing understanding of how early vaccine-induced antibody repertoires relate to subsequent viral evolution.

**One sentence summary:** Antibody epitopes on SARS-CoV-2 spike determine prospective breadth and vulnerability to viral evolution.

## Introduction

Neutralizing antibodies targeting the severe acute respiratory syndrome coronavirus 2 (SARS-CoV-2) spike glycoprotein are a central component of protective immunity induced by infection and vaccination (*1, 2*). Since the start of the coronavirus disease 2019 (COVID-19) pandemic, extensive work has focused on defining the antigenic landscape of the spike protein and on determining how antibody responses initially confer protection but also exert immune pressure that ultimately leads to viral escape through mutations in the targeted epitopes. These studies have established the receptor-binding domain (RBD) as the dominant target of potent neutralizing antibodies (*3–10*), largely because antibodies that interfere with the interaction between spike and its host receptor, angiotensin-converting enzyme 2 (ACE2), can efficiently block viral entry. Conversely, select studies implicate RBD-directed antibodies with antibody-dependent enhancement of infection (*11, 12*). In parallel, the N-terminal domain (NTD) of the spike protein has emerged as a secondary but highly immunogenic target, harboring distinct epitopes and an antigenic supersite that is frequently targeted during early antibody responses (*13–15*). The recognized importance of NTD-directed immunity has prompted the formulation of updated vaccines based on fusion constructs of the RBD and NTD (e.g., mRNA-1283) (*16*).

Despite the potency of antibodies directed against these epitopes, the continued evolution of SARS-CoV-2 has revealed important limitations in immunodominant antibody responses. Indeed, successive variants of concern (VOCs) have accumulated mutations that selectively evade neutralization by RBD-and NTD-directed antibodies, even in the presence of robust serological responses, indicating that most conserved spike epitopes are not neutralizing (*17–19*). Structural and functional studies have shown that many RBD-directed, neutralizing antibodies rely on interactions within the ACE2 footprint, also known as the receptor-binding motif (RBM). In contrast, NTD-directed antibodies often engage flexible loops that tolerate insertions, deletions, and conformational rearrangements (*20–25*). These observations suggest that antibody effectiveness is shaped not only by epitope accessibility and neutralization potency, but also by the evolutionary plasticity of the targeted epitopic region. However, how different modes of spike recognition affect antibody breadth, durability, and mechanisms of escape remains to be further explored.

Plasmablast responses elicited following first exposure to SARS-CoV-2 antigens provide a unique window into the initial humoral immune landscape against this virus. Antibodies derived from these responses frequently utilize either germline sequences or their minimally mutated versions and, therefore, offer insight into the intrinsic structural solutions favored by the human immune system (*7, 26, 27*). While many such antibodies have been isolated and characterized individually, few studies have systematically integrated high-resolution structural information with functional assays to understand how epitope specificity, binding geometry, and spike conformational dynamics collectively shape neutralization outcomes and susceptibility to viral evolution.

In a previous study, we described monoclonal antibodies (mAbs) elicited by mRNA vaccination originating from early plasmablast B-cell responses (18). We noted a balanced epitopic distribution across the spike glycoprotein, with mAbs engaging the RBD, NTD, and S2 domains. We identified seven neutralizing mAbs, all of which targeted either the RBD or the NTD. We also previously showed that these antibodies confer robust *in vivo* protection (*19*). We report here, high–resolution cryo–electron microscopy (cryo-EM) structures of all seven neutralizing mAbs bound to the spike trimer. By comparing antibodies that recognize canonical immunodominant epitopes with those that target less-explored regions of the spike, we sought to determine how distinct modes of antigen recognition influence prospective binding breadth, neutralization mechanisms, and susceptibility to antigenic drift. Our findings provide a framework for understanding why certain antibody responses predominate early yet rapidly lose effectiveness, whereas others may confer greater binding resilience in the face of ongoing viral evolution.

## Results

### Anti-RBD antibodies target Class I and Class II neutralizing epitopes

We isolated two RBD-directed neutralizing antibodies, PVI.V3-9 and PVI.V6-4, from early plasmablast responses after mRNA vaccination (18). To define the epitopes of two distinct RBD-targeting antibodies, we determined cryo-EM structures of Fab PVI.V3-9 and Fab PVI.V6-4 in complex with ancestral SARS-CoV-2 spike (USA/WA1/2020 strain, here referred as WA1) to 3.9 Å and 4.0 Å resolution, respectively (Figure 1, Figure 1—figure supplement 1-2). V3-9 is classified as a class I RBD antibody, binding exclusively to RBDs in its “up” conformation. In contrast, V6-4 belongs to class II RBD antibodies that can engage the RBD in both “up” and “down” configurations. In our cryo-EM reconstruction, the spike trimer has one RBD in the down conformation and two RBDs in the up conformation; Fab V6-4 can engage all three RBDs simultaneously, while Fab V3-9 occupies only the two RBDs in the up conformation.

**Figure 1.**
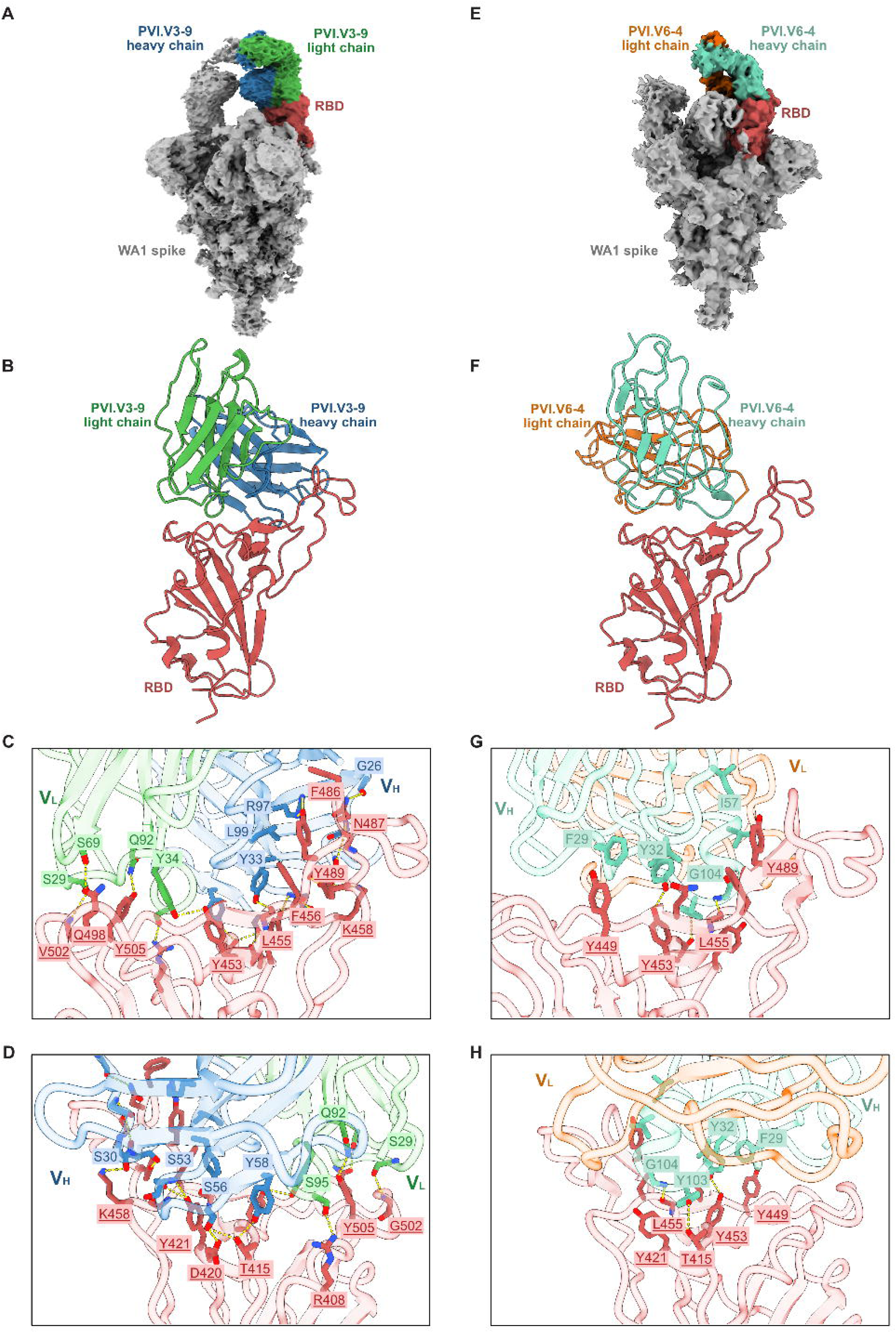
Anti-RBD antibodies recognizing Class I and Class II epitopes. (**A**) Cryo-EM reconstruction for spike-Fab V3-9 complex. (**B**) Fab V3-9 targets an epitope which overlaps with the ACE2 binding site on RBD and belongs to Class I. (**C**) Expanded view highlights interactions of V3-9 with RBD and illustrates hydrogen bonding and hydrophobic contacts of the interface (**D**) rotated 180° along the y-axis relative to C. (**E**) Cryo-EM reconstruction for spike-Fab V6-4 complex. (**F**) Fab V6-4 belongs to Class II and its epitope overlaps with the ACE2 binding site on the RBD. (**G**) Expanded view highlights interactions of V6-4 with RBD and illustrates hydrogen bonding and hydrophobic contacts of the interface (**H**) rotated 180° along the y-axis relative to G. Spike is colored grey, RBD red, V3-9 light chain green, V3-9 heavy chain blue, V6-4 light chain orange and V6-4 heavy chain teal.

V3-9, encoded by IGHV3-53 gene, is a class I neutralizing antibody that recognizes primarily the receptor-binding motif (RBM) (Figure 1A-B). V3-9 Fab thus binds in direct competition with ACE2 at the RBM, resulting in a total buried surface area of 1194.8 Å² at the interface. Multiple polar and non-polar contacts stabilize the RBD-antibody interface. Fifteen hydrogen bonds are formed between key RBD residues, including T415, D420, Y421, L455, R457, N460, Y473, A475, N487, and Y489, and heavy-chain complementarity-determining regions (HCDR1-3). Hydrophobic residues at the interface, including F456, K458, and F486, further reinforce the interaction through non-polar packing. An additional seven hydrogen bonds form between RBD residues R403, R408, and Y453 and the antibody light chain (CDRL1-3), involving S29, Y34, S69, Q92, Y93, and S95 (Figure 1C-D).

mAb V6-4 is an IGHV1-69 class II antibody that also targets the RBM on RBD and directly competes for binding with ACE2 (Figure 1E-F). The antigen-antibody interface buries a total surface area of 736 Å². Notably, the heavy chain alone mediates the interaction with the RBM, primarily through HCDR1 and HCDR3. Key contacts are established between RBD residues T415, Y453, L455, and N501 with CDRH residues G26, Y32, Y103, and G104. In addition, numerous hydrophobic interactions involving Y421, Y449, F456, Y489, and Q493 contribute to the stabilization of the complex (Figure 1G-H).

### Anti-NTD antibodies define two distinct neutralizing epitopes

We next characterized five NTD-targeting neutralizing antibodies isolated from the same early plasmablast response and determined four new structures and included one previously reported structure for detailed comparison. Among the five antibodies, three - PVI.V5-6, V6-7, and V6-2 are “NTD supersite binders,” engaging the supersite (site I) and aligning parallel to the spike trimer’s threefold axis. The remaining two antibodies-PVI.V6-11 and V6-14 (previously published; PDB ID: 7RBU) are “NTD lateral hydrophobic pocket binders”, bind outside the supersite on the lateral surface of the NTD and orient perpendicular to the trimer threefold axis. Together, these structures define two distinct modes of NTD recognition (*28*).

### NTD supersite binders

We determined the cryo-EM structure of the V5-6 Fab in complex with the SARS-CoV-2 spike trimer at a nominal local resolution of 3.76 Å. In our structure, all NTDs are occupied by Fab, consistent with full epitope engagement (Figure 2—figure supplement 1). The V5-6-spike complex reveals that antigen recognition is mediated by both antibody chains, with the heavy chain contributing a buried surface area of 682 Å² and the light chain of 345 Å² at the interface, underscoring substantial involvement of both chains in epitope engagement (Figure 2A-B). The heavy chain, encoded by IGHV3-7*01, establishes key contacts via Q14, R158, S161, and S162, where HCDR3 residues N101 and S105 make additional contacts with R158 and R246, and S31 of HCDR1 forms a hydrogen bond with Q14 (Figure 2C-D). Our structural analysis revealed that the light chain of V5-6, encoded by IGLV3-10*01, interacts predominantly with NTD residues Y144, H146, K147, N148, and R246. LCDR1 residues P28, K29, Y31, and Y33 form hydrogen bonds with H146, Y144, and N148, while D50 from LCDR2 contacts K147. Additionally, the Fab establishes contacts with N-linked glycans at N122, sandwiched between its HCDR2 and LCDR3, which may contribute to binding specificity or stabilization of the antigen-antibody interface (Figure 2—figure supplement 4A).

**Figure 2.**
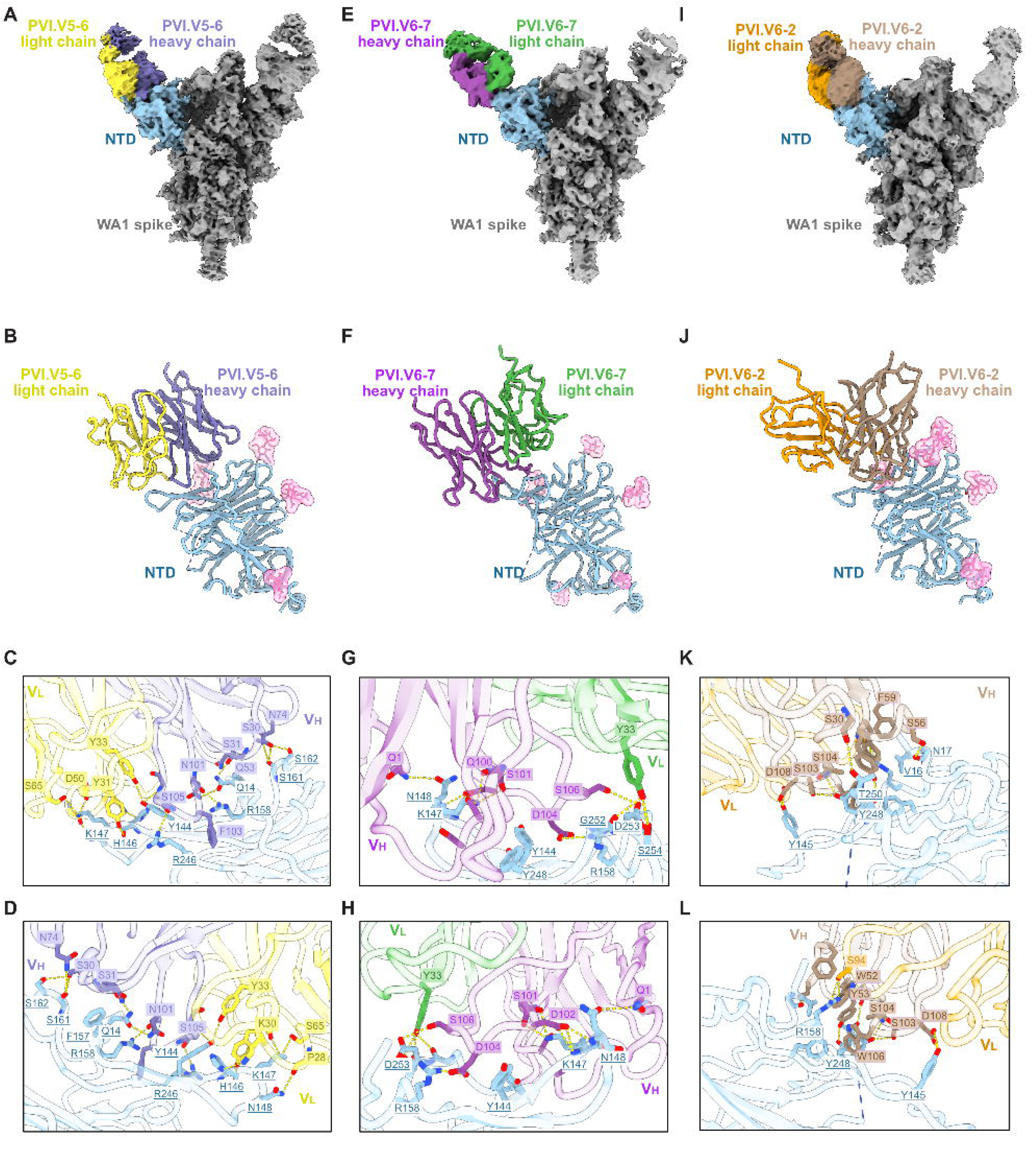
NTD binding antibodies recognize NTD antegenic supersite. **(A**) Cryo-EM reconstruction for spike-Fab V5-6 complex. (**B**) Fab V5-6 binds the supersite epitope on NTD. (**C**) Expanded view highlights interactions of V5-6 with NTD and illustrates hydrogen bonding and hydrophobic contacts at the interface (**D**) rotated 180° along the y-axis relative to C. (**E**) Cryo-EM reconstruction for spike-Fab V6-7 complex. (**F**) Fab V6-7 binds the supersite epitope on NTD. (**G**) Expanded view highlights interactions of V6-7 with NTD and illustrates hydrogen bonding and hydrophobic contacts at the interface (**H**) rotated 180° along the y-axis relative to G. (**I**) Cryo-EM reconstruction for spike-Fab V6-2 complex. (**J**) Fab V6-2 binds the supersite epitope on NTD. (**K**) Expanded view highlights interactions of V6-2 with NTD and illustrates hydrogen bonding and hydrophobic contacts at the interface (**L**) rotated 180° along the y-axis relative to K. Spike is colored grey, NTD light blue, V5-6 light chain yellow, V5-6 heavy chain indigo, V6-7 light chain green and V6-7 heavy chain purple, V6-2 light chain mustard, V6-2 heavy chain brown.

Similarly, we determined a cryo-EM structure of V6-7 Fab in complex with the WA1 spike trimer at a local resolution of 4.3 Å (Figure 2—figure supplement 2). V6-7 also targets the antigenic supersite (Figure 2E-F). The binding interface comprises an extensive network of polar and non-polar contacts. The heavy chain, encoded by IGHV4-39*01, dominates the interaction, contributing a buried surface area of 773 Å², while the light chain, derived from IGLV2-14*03, accounts for 115 Å². IGHV4-39 is among the most frequently utilized germline genes for NTD supersite-binding antibodies and is recognized as a public clonotype (*29*). The IGHV4-39/IGLV2-14 combination is among the most frequently observed antibody gene pairings in the human response to the SARS-CoV-2 spike, as reported in immune profiling studies (*30*). HCDR3 residues Q100, S101, D102, D104, and S106 engage multiple NTD residues, including H146, K147, N148, R158, and D253, while S33 of HCDR1 contributes an additional hydrogen bond with K147. The light chain, Y33 in LCDR1 further stabilizes the interface by contacting NTD residues G252, D253, and S254, (Figure 2G-H).

Finally, we determined the structure of V6-2 Fab complexed with WA1 spike extending to 4.4 Å resolution at the local antigen-combining interface (Figure 2—figure supplement 3). V6-2 also binds the NTD antigenic supersite (Figure 2 I-J). The heavy chain of V6-2 is encoded by IGHV3-33*01, which is very similar to the IGHV3-30 germline family and is one of the most frequently used IGHV genes among SARS-CoV-2 neutralizing antibodies (*31, 32*), while the light chain of V6-2 uses IGLV3-21*03. Structural analysis shows that antigen engagement is mediated by both antibody chains, with the heavy chain contributing a buried surface area of 730 Å² and the light chain accounting for 180 Å² at the interface, further supporting a dominant role for the heavy chain in epitope recognition. Detailed interaction mapping of the V6-2 Fab-NTD interface reveals a rich network of hydrogen bonds and hydrophobic contacts. On the heavy chain, N17, Y145, and T250 of the NTD form hydrogen bonds with S30 of HCDR1, Y53 and S56 of HCDR2, as well as S104 and D108 of HCDR3. In the light chain, S94 of LCDR3 establishes a hydrogen bond with R158 of the NTD. The N17 glycan also contacts HCDR2, further stabilizing the antigen-antibody interface (Figure 2—figure supplement 4B). In addition to these polar contacts, the interface features many hydrophobic interactions, such as those between W106, F59, and W52 of the heavy chain and F140 and V16 of the NTD, that strengthen the antigen-antibody binding surface (Figure 2K-L).

### NTD lateral hydrophobic pocket binders

We determined the cryo-EM structure of PVI.V6-11 at a local resolution of 4.53 Å. Structural analysis revealed that the spike trimer dissociated upon Fab V6-11 binding, resulting in a reconstruction with one Fab V6-11 engaged per NTD (Figure 3—figure supplement 1). For comparative purposes, we also analyzed PVI.V6-14, which we previously described (*28*). Both Fabs bind the lateral hydrophobic pocket of the NTD, previously shown to harbor a heme catabolite biliverdin; however, the mAbs approach the epitope at distinct angles (Figure 3A-H). V6-11 heavy chain is derived from IGHV1-46*01 and light chain from IGKV1-9*01, with calculated buried surface areas of 1025 Å² and 113 Å² for heavy and light chains, respectively. The Fab V6-11 paratope enters deeply into the NTD hydrophobic cavity via its HCDR3 loop, mediating both hydrophilic and hydrophobic contacts (Figure 3B-D). Specifically, W100, Q101, and W102 from HCDR3 interact hydrophobically with F175, H207, and F192, respectively. Additional polar contacts are established with T28 in HCDR1 which hydrogen bonds with P225, Y51 in HCDR2 with E224, and W100 in HCDR3 engaging Q173 (Figure 3C-D).

**Figure 3.**
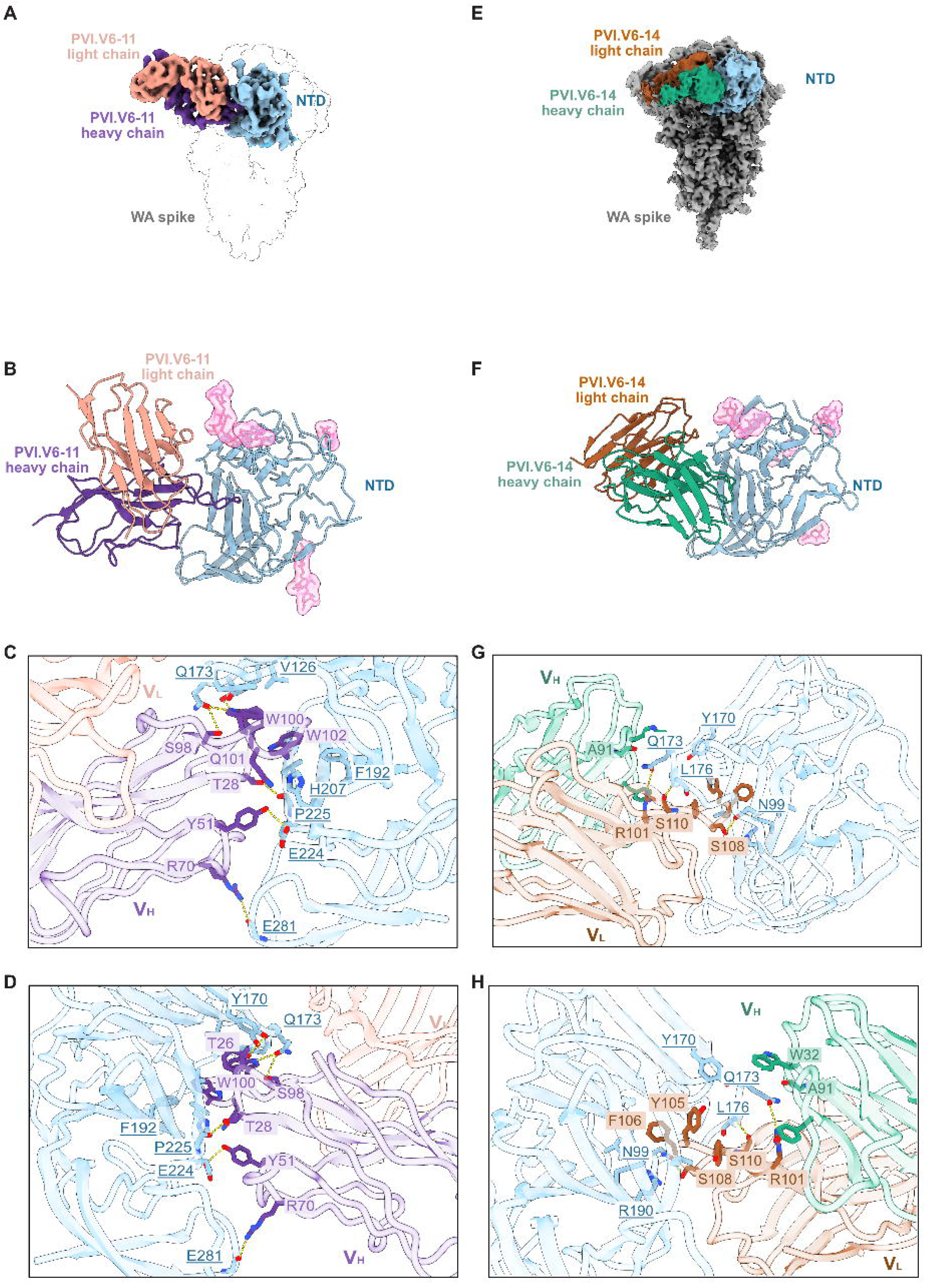
NTD lateral side binding antibodies recognize hydrophobic cavity. (**A**) Cryo-EM reconstruction for NTD-Fab V6-11 complex with spike trimer. As V6-11 dissociates the trimer, the focused map is shown in colors, and the putative trimer complex is shown in white (PDB:6VXX) to illustrate the incompatibility of the V6-11 complex with the trimeric spike. (**B**) Fab V6-11 targets the hydrophobic cavity of NTD lateral side. (**C**) Expanded view highlights interactions of V6-11 with NTD and illustrates hydrogen bonding and hydrophobic contacts at the interface (**D**) rotated 180° along the y-axis relative to C. (**E**) Cryo-EM reconstruction for spike-Fab V6-14 complex. (**F**) Fab V6-14 targets the hydrophobic cavity of NTD lateral side. (**G**) Expanded view highlights interactions of V6-14 with NTD and illustrates hydrogen bonding and hydrophobic contacts at the interface (**H**) rotated 180° along the y-axis relative to G. Spike is colored grey, NTD light blue, V6-11 light chain salmon, V6-11 heavy chain lavender, V6-14 light chain brown and V6-14 heavy chain green.

Fab V6-14 detailed cryo-EM structure is discussed in our previous paper (Figure 3E) (*28*). It features a heavy chain encoded by IGHV4-39*01 and a light chain encoded by IGKV1-12*01, with buried surface areas of 776 Å² and 147 Å², respectively. V6-14 also targets the NTD hydrophobic cavity using its HCDR3 residues, with Y104, Y105, and F106 contacting L226, Y170, R190, and F192 of NTD. The light chain further reinforces the interface through contacts by W32 in LCDR1 with P174 on NTD (Figure 3 G-H).

As observed in our cryo-EM structure, the spike trimer dissociation occurred upon V6-11 Fab binding to the NTD, and not with V6-14. To further investigate the structural basis for this dissociation, we superposed our NTD-Fab complex onto the SARS-CoV-2 spike trimer structure (PDB 6VXX). Strikingly, it revealed that Fab V6-11, when bound to the NTD within a full spike trimer, significantly clashed with the RBD of the neighboring S1 protomer, which likely causes the spike trimer to dissociate, as observed in our cryo-EM structure (Figure 3—figure supplement 2A). Conversely, Fab V6-14 does not clash with the adjacent RBD, consistent with our structural data in which all protomers remain intact (Figure 3—figure supplement 2B).

We have reported previously that V6-14 competes with biliverdin for binding in hydrophobic cavity of NTD (*28*). We superimposed V6-11 Fab bound NTD structure with biliverdin-NTD complex (*33*). Interestingly, we observed that V6-11 also overlaps with biliverdin’s binding footprint in hydrophobic cavity through its HCDR3 loop. It shares the same contact residues within NTD as V6-14 and biliverdin. N99, N121, R102, W104, R190, H207, L226 of NTD makes hydrophobic contacts with H99, W100, Q101, W102, L103 and D107 of HCDR3 (Figure 3— figure supplement 3).

### Prospective binding breadth of early plasmablast-derived neutralizing antibodies

We further characterized the binding breadth of these antibodies by measuring the binding across a panel of SARS-CoV-2 spike variants to determine the extent of cross-reactivity and structural tolerance to antigenic variation. To this end, we developed a high-throughput cell-based flow cytometry binding assay in which spike variants were expressed on the surface of HEK293T cells. Antibody binding was then quantified by flow cytometry and represented as a heatmap in Figure 4A with the gating strategy shown in Figure 4—figure supplement 1. Out of the two RBD targeting mAbs, V3-9 maintained binding to spike from WA1, B.1.1.7 (Alpha), P.1 (Gamma), and B.1.617.2 (Delta), but lost reactivity against Omicron and all subsequent variants (BA.1, BQ.1.1, BA.5, BA.2.86, XBB.1.5, and JN.1). In contrast, antibody V6-4 retained binding across pre-Omicron and early Omicron lineages, along with BA.2.86, with reduced binding to BA.1, BQ.1.1 and BA2.86.

**Figure 4.**
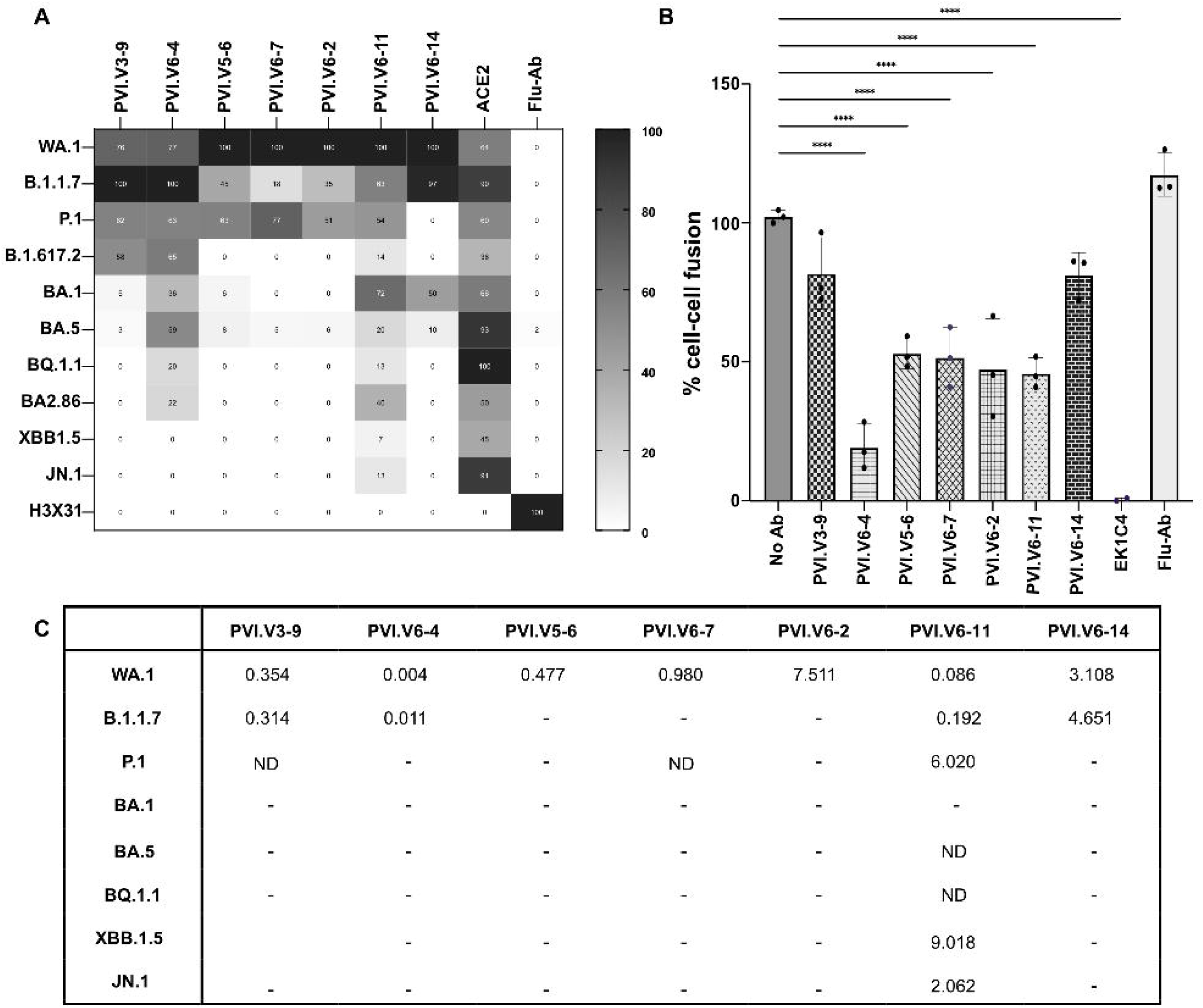
Functional characterization of SARS-CoV-2 spike-targeting antibodies. **(A)** Heatmap of relative binding to SARS-CoV-2 spike variants by flow cytometry. Percentage of spike positive cells binding each mAb was normalized such that the maximum % binding observed for each antibody across all variants is set to 100%. ACE2 and Flu-Ab binding to spike and variants are taken as positive and negative controls. All the SARS-CoV-2 spike variants are arranged as per chronological emergence. Data are representative of 2 independent biological repeats. **(B)** Cell fusion assay-Quantification of syncytia formation of WA1 spike. Cell-cell fusion was represented as GFP positive area which was normalized to spike with no Ab. Shown are the means and standard deviation (SD) of three replicates. EK1C4 and Flu-Ab are positive and negative control for cell fusion inhibition respectively. Data are representative of 2 independent biological repeats. Statistical significance was determined by one-way ANOVA followed by Tukey’s multiple-comparisons test (α = 0.05). ****P < 0.0001 **(C)** Table representing K_D_ values. K_D_ values for the Fab binding to spike trimer are represented in µM. “-” represents no measurable binding under the assay conditions. ‘ND=not determined’ represents binding detected but saturation was not reached within the tested concentration range.

The mAb completely lost the binding to the more recent XBB.1.5 and JN.1 subvariants. All three NTD supersite binders exhibited cross-reactivity with B.1.1.7 and P.1 but lost binding to B.1.617.2 and all tested Omicron subvariants. Out of the two NTD hydrophobic cavity binders, V6-11 showed cross-reactivity to all tested subvariants with lower binding to B.1.617.2, BQ.1.1, BA.4, XBB.1.5, and JN.1 while V6-14 cross-reacted with B.1.1.7 and BA.1 with little or no binding with BA.4 subvariant.

To further quantify the binding properties of these antibodies, we measured the binding affinities of Fab fragments to purified SARS-CoV-2 spike trimers by ELISA. The corresponding K_D_ values are summarized in Figure 4C, with the binding curves shown in Figure 4—figure supplement 2. PVI.V6-4 showed the strongest measurable binding to the ancestral WA.1 and B.1.1.7 spikes, with K_D_ values of 0.004 and 0.011 µM, respectively. In contrast, PVI.V6-11 displayed the broadest binding profile, with measurable binding to WA.1, B.1.1.7, P.1, BA.5, BQ.1.1, XBB.1.5, and JN.1. The remaining antibodies showed more restricted binding, with measurable binding primarily to WA.1 and, for some antibodies, P.1, and no measurable binding to most of the Omicron variants tested. Notably, the ELISA measurements are different from the cell-surface binding profiles observed by FACS, as PVI.V6-11 showed binding to BA.1 and BA.2.86 by FACS but no measurable Fab binding to these variants by ELISA. These differences likely reflect differences in antigen presentation, assay format, and antibody valency between the two approaches.

To investigate the structural basis of cross-reactivity of our antibodies, we performed sequence analysis of the RBDs from all tested SARS-CoV-2 variants, focusing on interface residues engaged by anti-RBD antibodies. The loss of binding of V3-9 to Omicron variants is attributed to the Q498R and Y505H mutations, which disrupt key interactions with V3-9. V6-4 loses binding to XBB.1.5 and JN.1 due to L455F and L455S mutation, respectively (Figure 4—figure supplement 3).

Similarly, to understand why all NTD top binders in our panel lose binding to Delta and Omicron variants, we conducted sequence analysis of the NTD of these variants, comparing the interface residues involved in Fab recognition (Figure 4—figure supplement 4). We found that mutations and deletions within the N2, N3, and N5 loops, critical components of the NTD antigenic supersite, correlate with the observed loss of binding of the supersite-directed antibodies. Although the N2 loop does not directly contact the Fab in our structures, mutations in this loop may indirectly influence antibody recognition by altering the local architecture and conformational flexibility of NTD antigenic supersite. Comparative structural analysis of our WA1 NTD-Fab complexes with published structures of XBB1.5 and BA.5 spike complexes with NTD-directed antibodies further reveals substantial loop flexibility in these regions. The N1-N5 loops are disordered in most reported structures, but some of these loops become ordered when complexed with mAbs (*14*). Conformational flexibility and structural rearrangements of the N2 loop (67-79 aa), N3 loop (141-156 aa), and N5 loop (246-260 aa) in the NTD may contribute to the observed loss of binding of our antibodies against Delta, Omicron, and other emerging variants. These loops constitute a critical component of the NTD antigenic supersite and interact extensively with all three antibodies studied (V5-6, V6-7, and V6-2). The dynamic nature of these NTD loops could create steric clashes with antibody paratopes, thereby diminishing antibody accessibility and reducing binding (Figure 4—figure supplement 5). Many structural studies have implicated NTD loop mobility as a key mechanism for immune escape in emerging SARS-CoV-2 variants (*13, 14, 34, 35*). Sequence analysis of the NTD across all tested variants shows that most residues forming the hydrophobic pocket contacted by V6-11 and V6-14 are conserved. Structural comparison of the WA1 NTD-

Fab complexes indicates that, although both Fabs target largely overlapping residues, V6-11 buries a larger surface area (1138 Å²) and makes more hydrogen bonds to conserved cavity residues, whereas V6-14, with a surface area of 923 Å², depends only on hydrophobic contacts. These differences suggest that loop remodeling in P.1 and BA.2.86 VOCs might disrupt V6-14 binding, while the deeper engagement of V6-11 tolerates such changes and supports its broader binding profile (Figure 4—figure supplement 6).

### Cell fusion inhibition potency

Given our cryo-EM structures, we propose that the RBD- and NTD top-binding mAbs neutralize by either direct competition with ACE2 or steric hindrance, respectively, but the neutralization mechanism of the hydrophobic pocket-targeting mAbs remains unclear. We, therefore, performed a cell-cell fusion inhibition assay. Syncytia formation, a hallmark of SARS-CoV-2 infection and lung pathogenesis, is driven by interaction between the viral spike protein and the host ACE2 receptor (*36*). To investigate whether our antibodies could inhibit viral spike-mediated cell fusion, we used a split-GFP syncytia assay in VeroE6-TMPRSS2 reporter cells, in which the two complementary GFP halves are present in separate cell populations. Both cells express ACE2 and TMPRSS2. The cells were mixed and transfected with the SARS-CoV-2 WA1 spike. We incubated the antibody for 24 h and quantified syncytia formation by measuring the total GFP-positive area per well using the Celigo imaging system. We observed that V6-4 inhibited cell-cell fusion by 80%, whereas V3-9 inhibition reached only 20%. All NTD top binders showed approximately 50% inhibition of cell-cell fusion, while the NTD hydrophobic pocket binding V6-11 also showed 50% inhibition, and V6-14 inhibited fusion by 20% (Figure 4B). Statistical analysis revealed significant inhibition for V6-4, V5-6, V6-7, V6-2, and V6-11 whereas the inhibition observed for V3-9 and V6-14 did not reach statistical significance.

## Discussion

The complex interplay among epitope accessibility, conformational dynamics of the spike glycoprotein, and evolutionary constraints imposed by viral fitness shapes neutralizing antibody responses against SARS-CoV-2. Previous work has established the RBD as the dominant target of potent neutralizing antibodies (*5, 8, 10, 18, 37*), while the NTD has emerged as a secondary but highly immunogenic site that is particularly prone to immune escape, as it tolerates insertions and deletions in its flexible loops (*13, 28, 38, 39*). Plasmablast-derived antibodies represent a snapshot of early vaccine-induced humoral response. Although later affinity maturation and clonal selection may preserve or alter these binding modes in serum, our study defines the starting point from which subsequent antibody evolution can be understood. By structurally and functionally characterizing early plasmablast–derived antibodies targeting both domains, we provide here mechanistic insights into how distinct modes of spike recognition translate into differences in prospective binding breadth, durability, and susceptibility to antigenic drift.

Previous studies have shown that class I RBD antibodies, encoded by public germline genes such as IGHV3-53/IGHV3-66, predominate in early SARS-CoV-2 antibody responses and potently neutralize ancestral viruses through direct competition with the host receptor, ACE2 (*27, 40–42*). Although we have shown that rare members of this class of antibodies retain robust neutralizing capacity (*42*) despite rapid viral evolution, most lose neutralizing efficacy against Omicron and later variants. Our current data provide a structural rationale for this phenomenon; by engaging a highly constrained receptor-binding motif in a conformation-dependent manner, class I antibodies impose strong selective pressure on residues that are simultaneously dispensable for ACE2 binding. The emergence of Omicron-associated mutations, such as those at positions Q498 and Y505 (*35, 43*) exemplifies how substitutions at the periphery of the ACE2 interface can abrogate antibody binding while preserving ACE2 binding and, consequently, viral entry. These observations help explain the recurrent escape from this antibody class observed across multiple independent studies.

In contrast, class II RBD antibodies exhibit greater tolerance to antigenic variation by binding the RBD in both “up” and “down” conformations and by engaging less variable RBM residues (*10, 44, 45*). Our structural and binding analyses support this model, while also revealing its limitations. Indeed, although class II recognition may endow the antibodies with broader binding breadth, including the early Omicron lineages, mutations at key hydrophobic residue positions such as L455 ultimately lead to the loss of binding. These observations suggest that while conformational flexibility may afford greater resistance to immune escape, it does not eliminate the fundamental constraint imposed by targeting a region of the spike that is under intense immune and evolutionary pressure (*25, 46, 47*).

Early in the COVID-19 pandemic, multiple groups identified the NTD antigenic supersite as a major target of neutralizing antibodies, often encoded by recurrent germline genes and capable of potent neutralization in vitro (*13, 14, 28, 38, 39, 48, 49*). Yet, most NTD-directed antibodies were among the first to lose activity as variants of concern emerged. Our work reinforces and mechanistically extends these observations by demonstrating that antibodies recognizing the NTD supersite converge on highly flexible loop regions that readily accommodate deletions, insertions, and conformational rearrangements (*45, 50–52*). Structural comparisons across variants suggest that loop plasticity, rather than single-point mutations, is the dominant mechanism of escape, enabling major structural remodeling of the supersite without compromising spike function. This plasticity likely explains why NTD supersite antibodies are repeatedly elicited yet consistently short-lived in their protective capacity.

Importantly, our study also highlights an alternative mode of NTD recognition that challenges the prevailing view of the NTD as an inherently poor epitopic region for broadly neutralizing antibodies. Antibodies targeting the lateral hydrophobic cavity of the NTD engage a structurally conserved pocket that is functionally constrained by its role in binding small molecules such as biliverdin (*28, 33*). The breadth exhibited by these antibodies across diverse SARS-CoV-2 variants suggests that this site is less permissive to viral escape. We also speculate that certain antibodies of the same class may destabilize the spike trimer and thus neutralize the virus via a mechanism that is distinct from the classical ACE2 blockade, whereby perturbation of spike architecture or conformation impairs membrane fusion. These findings suggest that effective neutralization does not require direct interference with ACE2 binding. Instead, antibodies that exploit structurally constrained allosteric sites may achieve broader activity by targeting epitopes that are less permissive to evolution under immune pressure. Such mechanisms may be particularly relevant for limiting cell–cell fusion and syncytia formation, which have been increasingly recognized as contributors to SARS-CoV-2 pathogenesis and viral spread *in vivo*.

Taken together, our antibody panel reveals two recognition strategies for RBD-directed antibodies, which target the ACE2-binding surface through distinct class I and class II binding sites, and two strategies for NTD-directed antibodies, which either recognize the flexible supersite with different angles of approach or bind the conserved lateral hydrophobic pocket. While early antibody responses favor highly accessible and potently neutralizing epitopes on the RBD and NTD supersite, these targets are also among the most mutable under immune pressure. In contrast, antibodies targeting structurally conserved, functionally constrained regions of the spike, such as the NTD hydrophobic cavity, may represent less immunodominant but more evolutionarily stable targets in the face of viral evolution. These insights support vaccine and immunogen design strategies that direct humoral immunity toward conserved, immune-subdominant epitopes that can complement the canonical neutralizing epitopes on the RBD and NTD and achieve broader, more durable protection against both current and future sarbecoviruses.

## Materials and Methods

### Cell-based binding assay by flow cytometry

The plasmid expressing the full-length SARS-CoV-2 variant spike protein were transfected with human embryonic kidney 293T cells (HEK293T) at a ratio of 1:3 using polyethylenimine (PEI) in Optimized Minimal Essential Medium (Opti-MEM, Gibco) transfection medium. After 5 h of incubation at 37 °C with 5% CO2, the cells were supplemented with Dulbecco’s modified Eagle’s medium (DMEM, Gibco) containing of 10% Fetal Bovine Serum (FBS, corning). After 72 h, the cells were harvested and stained with test antibodies at 10 ug/ml at 4°C for 1 h. The cells were washed with FACS buffer (1x PBS (phosphate buffered saline) containing 1% BSA and 1mM EDTA), then PE (phycoerythrin)-conjugated goat anti-human IgG Fc antibody (Invitrogen-12-4998-82) was added to the cells at a ratio of 1:250 and incubated at 4 °C for 30 min. Finally, cells were resuspended, and binding of antibody was quantified by AttuneNxT, (Invitrogen) and data was analyzed by flowJo.

### ELISA

Binding of Fab antibodies to SARS-CoV-2 spike trimers and their variants was assessed by ELISA. Purified spike trimers were coated onto ELISA plates at 2 μg/mL and incubated for antigen immobilization overnight at 4C. Plates were blocked with 3% skim milk in PBST for 2 h and subsequently washed twice with 1x PBS containing Tween-20 (PBST). Fab antibodies were serially diluted fivefold (1:5) and added to the coated plates, followed by incubation for 1 h. Plates were washed twice with PBST and incubated with anti-human IgG Fab-specific secondary antibody (Sigma-Aldrich-A0293) for 1 h. Plates were then washed twice with PBST, and bound antibody was detected using 3,3′,5,5′-tetramethylbenzidine (TMB) substrate. Absorbance was measured following substrate development. Binding curves were generated by plotting the ELISA absorbance against Fab concentration and analyzed by nonlinear regression using the “One site-specific binding” model in GraphPad Prism (v10.4.2).

### Cell-cell fusion assay

This assay uses a split-GFP system comprising GFP1-10 and GFP11 fragments that emit fluorescence upon cell fusion (*53*). VeroE6-TMPRSS2 cells that stably expresses GFP 1-10, and GFP 11 fragments were used. Briefly, 40 ng of wild-type (WA1) spike plasmid per well (96-well plate) was complexed with TransIT-LT1 transfection reagent at a ratio of 1:6 (Takara; MIR2300) in 10ul of OptiMEM for reverse transfection. This transfection complex was then combined with 15,000 cells from each VeroE6-TMPRSS2-HA-GFP1-10 and VeroE6-TMPRSS2-BSR-GFP11-flag expressing populations in a final volume of 90 µL of OptiMEM. The mixture was seeded into black 96-well plates (Greiner, 655090). After 6 hours of incubation to allow transfection, the media was replaced with DMEM supplemented with 10% FBS and antibody at 100 µg/mL. Cells were fixed 24 hours post-transfection with 4% paraformaldehyde in phosphate buffered saline (PBS) for 10 minutes. GFP fluorescence, reflecting cell-cell fusion events, was quantified using a Celigo Image Cytometer (Nexelcom Bioscience, Version 4.1.3.0). Fusion signals were normalized relative to spike expression.

### Spike, Fab, and IgG expression and purification

Expression and purification of stabilized SARS-CoV-2 6P ectodomain was conducted as previously described (*28*). Briefly, constructs encoding the SARS-CoV-2 S ectodomain with hexa-His tag were used to transiently transfect Expi293F cells (Gibco). Five days after transfection, supernatants were harvested, and spike proteins were purified by cobalt resin and size-exclusion chromatography. Peak fractions from size-exclusion chromatography were identified by Sodium Dodecyl Sulfate-Polyacrylamide Gel Electrophoresis (SDS-PAGE), and fractions corresponding to spike trimers were pooled and stored at 4°C. Fabs and IgGs were expressed, and purified as previously described (*28*).

### Cryo-electron microscopy sample preparation and data collection

The SARS-CoV-2 WA1 HexaPro spike was mixed with Fabs at a concentration of 2 mg/ml, using a 1.5 molar excess of Fab, and incubated for 20 min at room temperature. Samples were prepared on UltrAuFoil gold R1.2/1.3 grids (Ted Pella, Inc) and were subjected to glow discharging in a PELCO easiGlow device (air pressure: 0.39 mBar, current: 28 mA, duration: 25 s) before sample preparation. Immediately before grid preparation, fluorinated octyl-maltoside was added to the complex at a final concentration of 0.02% wt/vol. Next, 3 μl of sample were applied to grids, which were blotted for 6 s using grade 595 filter paper (Ted Pella Inc.) at a blot force of 0 at 22°C and 95% humidity. The samples were then plunge-frozen in liquid ethane using a Vitrobot Mark IV system (ThermoFisher Scientific). Imaging was conducted on a ThermoFisher Scientific Titan Krios G3i transmission electron microscope (300 kV, XFEG electron source) featuring a three-condenser lens system. The microscope was equipped with Gatan BioQuantum energy filter (15 eV slit) and a Gatan K3 direct electron detector. Data acquisition was performed using Leginon v3.7.

### Cryo-electron microscopy data processing, structure modeling, and refinement

Data process workflow, including motion correction, Contrast Transfer Function (CTF) estimation, particle picking and extraction, 2D classification, ab initio reconstruction, homogeneous refinement, heterogeneous refinement, non-uniform refinement, local refinement and local resolution estimation, were carried out in cryoSPARCv.3.3.1 The overall resolution was 3.11 Å, 3.98 Å, 3.28 Å, 3.85 Å, 3.80 Å and 4.55 Å for the map of WA1 spike with V3-9, V6-4, V5-6, V6-7, V6-2 and V6-11 respectively. To resolve the RBD-antibody interface and NTD-antibody interface, local refinements were performed, a mask encompassing the RBD-antibody or NTD-antibody region was used for refinement. The two half-maps from this refinement were sharpened using DeepEMhancer v1.0 (*54*). The reported resolutions are based on the gold-standard Fourier shell correlation criterion of 0.143.

The focused maps sharpened with DeepEMhancer were used for model building. The initial models was created with ABodyBuilder2 (*55*) and then manually refined with COOT v0.9.8.91 (*56*). N-linked glycans were added manually in COOT using the glyco extension. The model underwent further refinement in Phenix v1.21.2_5419, employing real-space refinement with Ramachandran and rotameric restraints, and was validated using MolProbity v4.4 (*57*) (Supplementary table 1). The structural biology software was compiled and made available through SBGrid (*58*). The overall workflow for each antibody data processing is elaborated in Figure 1—figure supplement 1-2, Figure 2—figure supplement 1-3, Figure 3—figure supplement 1.

## Acknowledgements

We acknowledge Adolfo García-Sastre and Alba Escalera for providing the split-GFP VeroE6-TMPRSS2 cell lines. We acknowledge support from the Irma T. Hirschl/Monique Weill-Caulier Trust (G.B.). Some of this work was performed at the National Center for CryoEM Access and Training (NCCAT) and the Simons Electron Microscopy Center located at the New York Structural Biology Center, supported by the NIH Common Fund Transformative High Resolution Cryo-Electron Microscopy program (U24 GM129539), and by grants from the Simons Foundation (SF349247) and NY State Assembly. This work was supported in part through the computational and data resources and staff expertise provided by Scientific Computing and Data at the Icahn School of Medicine at Mount Sinai and supported by the Clinical and Translational Science Awards (CTSA) grant UL1TR004419 from the National Center for Advancing Translational Sciences. Research reported in this publication was also supported by the Office of Research Infrastructure of the National Institutes of Health under award number S10OD026880 and S10OD030463. The content is solely the responsibility of the authors and does not necessarily represent the official views of the National Institutes of Health.

## Author contributions

Conceptualization: GB. Formal analysis: GB, DJ. Funding acquisition: AHE, GB. Investigation: CGA, DCA, GB, IAS, DJ. Methodology: CGA, GB, DJ. Project administration: GB. Resources: AHE, GB, FK, VS. Supervision: GB. Validation: GB, DJ. Visualization: DJ. Writing—original draft: GB and DJ. Writing—review and editing: AHE, GB, DJ, FK, VS.

## Data and materials availability

The EM maps have been deposited in the Electron Microscopy Data Bank (EMDB) under accession code EMD-75040, EMD-75041, EMD-75042, EMD-75043, EMD-75044, EMD-75045 for V3-9, V6-4, V5-6, V6-7, V6-2, V6-11 respectively and the accompanying atomic coordinates in the Protein Data Bank (PDB) under accession code 10BF (V3-9), 10BG (V6-4), 10BH (V5-6), 10BI (V6-7), 10BJ (V6-2), 10BK (V6-11). The content is solely the responsibility of the authors and does not necessarily represent the official views of the National Institutes of Health.

## Statistical analysis

Data were analyzed using GraphPad PRISM 10 for macOS (version 10.3.0).

## Figure supplements

**Figure 1 figure supplement 1. Overview of cryo-EM data processing and local resolution analysis.** (**A**) Representative cryo-electron micrograph of the SARS-CoV-2 WA spike V3-9 complex. (**B**) Selected 2D class averages of the WA1 spike ectodomain bound to V3-9. (**C**) Local resolution maps and gold-standard Fourier shell correlation (FSC) curves for the global spike-Fab reconstruction generated using cryoSPARC v3.3.1. (**D**) and locally refined receptor-binding domain (RBD)-Fab complex. (**E**) Schematic representation of the overall cryo-EM data processing workflow. (**F**) Orientation distribution plot (**G**) Cryo-EM density of interface residues.

**Figure 1 figure supplement 2. Overview of cryo-EM data processing and local resolution analysis.** (**A**) Representative cryo-electron micrograph of the SARS-CoV-2 WA1 spike V6-4 complex. (**B**) Selected 2D class averages of the WA1 spike ectodomain bound to V6-4. (**C**) Local resolution maps and gold-standard Fourier shell correlation (FSC) curves for the global spike-Fab reconstruction generated using cryoSPARC v3.3.1. (**D**) and locally refined receptor-binding domain (RBD)-Fab complex. (**E**) Schematic representation of the overall cryo-EM data processing workflow. (**F**) Orientation distribution plot (**G**) Cryo-EM density of interface residues.

**Figure 2 figure supplement 1. Overview of cryo-EM data processing and local resolution analysis.** (**A**) Representative cryo-electron micrograph of the SARS-CoV-2 WA1 spike V5-6 complex. (**B**) Selected 2D class averages of the WA1 spike ectodomain bound to V5-6. (**C**) Local resolution maps and gold-standard Fourier shell correlation (FSC) curves for the global spike-Fab reconstruction generated using cryoSPARC v3.3.1. (**D**) and locally refined N-terminal domain (NTD)-Fab complex. (**E**) Schematic representation of the overall cryo-EM data processing workflow. (**F**) Orientation distribution plot (**G**) Cryo-EM density of interface residues.

**Figure 2 figure supplement 2. Overview of cryo-EM data processing and local resolution analysis.** (**A**) Representative cryo-electron micrograph of the SARS-CoV-2 WA1 spike V6-7 complex. (**B**) Selected 2D class averages of the WA1 spike ectodomain bound to V6-7. (**C**) Local resolution maps and gold-standard Fourier shell correlation (FSC) curves for the global spike-Fab reconstruction generated using cryoSPARC v3.3.1. (**D**) and locally refined N-terminal domain (NTD)-Fab complex. (**E**) Schematic representation of the overall cryo-EM data processing workflow. (**F**) Orientation distribution plot (**G**) Cryo-EM density of interface residues.

**Figure 2 figure supplement 3. Overview of cryo-EM data processing and local resolution analysis.** (**A**) Representative cryo-electron micrograph of the SARS-CoV-2 WA1 spike V6-2 complex. (**B**) Selected 2D class averages of the WA1 spike ectodomain bound to V6-2. (**C**) Local resolution maps and gold-standard Fourier shell correlation (FSC) curves for the global spike-Fab reconstruction generated using cryoSPARC v3.3.1. (**D**) and locally refined N-terminal domain (NTD)-Fab complex. (**E**) Schematic representation of the overall cryo-EM data processing workflow. (**F**) Orientation distribution plot (**G**) Cryo-EM density of interface residues.

**Figure 2 figure supplement 4.** (**A**) N122 glycan of spike interacting with CDRH2 and CDRL3. (**B**) N17 glycan of spike interacting with CDRL3. Glycans are shown as sticks in pink and antibody interacting residues are shown as sticks.

**Figure 3figure supplement 1. Overview of cryo-EM data processing and local resolution analysis.** (**A**) Representative cryo-electron micrograph of the SARS-CoV-2 WA1 spike V6-11 complex. (**B**) Selected 2D class averages of the WA1 spike ectodomain bound to V6-11. (**C**) Local resolution maps and gold-standard Fourier shell correlation (FSC) curves for the local spike-Fab reconstruction generated using cryoSPARC v3.3.1. (**D**) Schematic representation of the overall cryo-EM data processing workflow. (**E**) Orientation distribution plot (**F**) Cryo-EM density of interface residues.

**Figure 3 figure supplement 2. Analysis of heavy chain clashes with spike protomers** (**A**) The heavy chain of PVI.V6-11 exhibits steric clashes with the RBD of protomer B.(**B**) the heavy chain of PVI.V6-14 does not display any clashes with protomer B of the spike. PVI.V6-11 heavy chain is shown in lavender and PVI.V6-11 light chain is shown in salmon, PVI.V6-14 heavy chain is shown in green and PVI.V6-14 light chain is shown in orange. Protomer A, B and C of spike is shown in light blue, tan and gray respectively.

**Figure 3 figure supplement 3. PVI.V6-11 competes with biliverdin.** Structural superposition of biliverdin bound NTD (PDB: 7B62) with PVI.V6-11 bound NTD (this study). Biliverdin is shown in green. PVI.V6-11 heavy chain is shown in lavender with CDRH3 loop highlighted in sticks. NTD is shown in light blue with interacting residues highlighted in sticks.

**Figure 4 figure supplement 1. Gating strategy used in cell-based flow cytometry binding assay**. In FSC-A versus FSC-H, identification of the singlets; in SSC-A versus SSC-H, identification of the singlets; in FSC-A versus SSC-A, all selected cells, in SSC-A versus PE-A, identification of positive cells.

**Figure 4 figure supplement 2. ELISA binding curves of Fab antibodies to SARS-CoV-2 spike trimer.** Binding profiles of Fabs against the indicated spike variants. Dotted lines represent nonlinear regression fits using the “One site-Specific binding” model in GraphPad Prism. Representative binding curves from one of two independent experiments are shown. The corresponding K_D_ values are summarized in Figure 4C.

**Figure 4 figure supplement 3. Multiple sequence alignment of RBD with epitope residues of RBD binders.** SARS-CoV-2 spike receptor-binding domain (RBD; amino acids 319-542) of WA1, B.1.1.7, P.1, B.1.617.2, BA.1, BQ.1.1, BA.5, BA.2.86, XBB.1.5, and JN.1. Epitope residues of PVI.V3-9, PVI.V6-4 and ACE2 are indicated in heat map. Blue represents the heavy chain, yellow represents the light chain, white represents no interaction and brown represents ACE2 interaction.

**Figure 4 figure supplement 4. Multiple sequence alignment of NTD with epitope residues of NTD supersite binders.** SARS-CoV-2 spike N-terminal domain (NTD; amino acids 14-306) of WA1, B.1.1.7, P.1, B.1.617.2, BA.1, BQ.1.1, BA.5, BA.2.86, XBB.1.5, and JN.1. Epitope residues of Fab PVI.V5-6, PVI.V6-7 and PVI.V6-2 are indicated in heat map. Blue represents the heavy chain, yellow represents the light chain, green represents both heavy and light chain interaction and white represents no interaction.

**Figure 4 figure supplement 5. Structural overlay of SARS-CoV-2 NTD variants reveals loop deviations and antibody clash.** (**A**) Superimposition of the NTD from the WA1 spike in complex with PVI.V5-6 onto NTD structures from BA.5 NTD with an RBD-bound antibody (PDB 8GTP), XBB.1.5 NTD (unliganded, PDB 8VKK), and XBB.1.5 in complex with an NTD-directed antibody (PDB 9CCJ) reveals significant conformational deviations in the N2, N3, and N5 loops. (**B**) Structural overlay shows that loop rearrangements in XBB.1.5 and BA.5 produce steric clashes with the PVI.V5-6 paratope. (**C**) Superimposition of the NTD from the WA1 spike in complex with PVI.V6-7 onto NTD structures from BA.5 NTD with an RBD-bound antibody (PDB), XBB.1.5 NTD (unliganded, PDB), and XBB.1.5 in complex with an NTD-directed antibody (PDB) reveals significant conformational deviations in the N2, N3, and N5 loops among these variants. (**D**) Structural overlay shows that loop rearrangements in XBB.1.5 and BA.5 produce steric clashes with the PVI.V6-7 paratope. (**E**) Superimposition of the NTD from the WA1 spike in complex with PVI.V6-7 onto NTD structures from BA.5 NTD with an RBD-bound antibody (PDB), XBB.1.5 NTD (unliganded, PDB), and XBB.1.5 in complex with an NTD-directed antibody (PDB) reveals significant conformational deviations in the N2, N3, and N5 loops among these variants. (**F**) Structural overlay shows that loop rearrangements in XBB.1.5 and BA.5 produce steric clashes with the PVI.V6-2 paratope.

**Figure 4 figure supplement 6. Multiple sequence alignment of NTD with epitope residues of NTD lateral hydrophobic pocket binders.** SARS-CoV-2 spike N-terminal domain (NTD; amino acids 14-306) of WA1, B.1.1.7, P.1, B.1.617.2, BA.1, BQ.1.1, BA.5, BA.2.86, XBB.1.5, and JN.1. Epitope residues of Fab PVI.V6-11, and PVI.V6-14 are indicated in heat map. Blue represents the heavy chain, yellow represents the light chain, green represents both heavy and light chain interaction and white represents no interaction.

**Supplementary table 1.** Cryo-EM data collection and model validation statistics.

|  | **PDB ID 10BF EMD-75040** | **PDB ID 10BG EMD-75041** | **PDB ID 10BH EMD-75042** | **PDB ID 10BI EMD-75043** | **PDB ID 10BJ EMD-75044** | **PDB ID 10BK EMD-75045** |
|---|---|---|---|---|---|---|
| Grid Type | UltrAuFoil gold R1.2/1.3 | UltrAuFoil gold R1.2/1.3 | UltrAuFoil gold R1.2/1.3 | UltrAuFoil gold R1.2/1.3 | UltrAuFoil gold R1.2/1.3 | Ultraufoil Gold R1.2/1.3 |
| Microscope/Voltage/Detector | Titan Krios/300 kV/Gatan K3 summit | Titan Krios/300 kV/Gatan K3 summit | Titan Krios/300 kV/Gatan K3 summit | Titan Krios/300 kV/Gatan K3 summit | Titan Krios/300 kV/Gatan K3 summit | Titan Krios/300 Kv/Gatan K3 Summit |
| Magnification | 105,000 | 105,000 | 105,000 | 105,000 | 105,000 | 105,000 |
| Recording Mode | counting | counting | counting | counting | counting | counting |
| Total Dose e-/Å<sup>2</sup>/s | 52.51 | 52.75 | 50.00 | 63.96 | 52.03 | 65.60 |
| Pixel Size (Å/pixel) | 1.069 | 1.076 | 1.058 | 1.083 | 1.076 | 1.076 |
| Defocus Range (µm) | 0.5 to 1.5 | 0.5 to 1.5 | 0.5 to 1.5 | 0.5 to 1.5 | 0.5 to 1.5 | 0.5 to 1.5 |
| No. Micrographs Used | 9,951 | 4,028 | 9,444 | 9,441 | 13,518 | 9,054 |
| Total Particles Picked | 1,471,147 | 1,489246 | 397,353 | 317,743 | 9,80,508 | 2,722,827 |
|  | **Model Validation** |  |  |  |  |  |
| Chains | 4 | 3 | 4 | 4 | 4 | 3 |
| Atoms | 3236 | 3356 | 4097 | 4091 | 4043 | 4100 |
| Residues | **Protein** |  |  |  |  |  |
|  | 416 | 430 | 504 | 513 | 505 | 506 |
|  | **Nucleotide** |  |  |  |  |  |
| Ligands | NAG: 2, BMA:1 | NAG: 1 | NAG: 7, BMA:1 | NAG: 7 | NAG: 9, BMA:1 | NAG: 6 BMA:2 |
|  | **Bonds (Rmsd)** |  |  |  |  |  |
| Length (Å) (# > 4sigma) | 0.003 (0) | 0.005 (0) | 0.005 (0) | 0.005 (0) | 0.007 (0) | 0.006 (0) |
| Angles (°) (# > 4sigma) | 0.803 (0) | 0.923 (4) | 0.825 (1) | 0.874 (0) | 1.514 (17) | 1.417 (9) |
| Molprobity Score | 1.49 | 1.45 | 1.54 | 1.38 | 1.73 | 1.29 |
| Clash Score | 4.27 | 1.98 | 3.36 | 2.62 | 4.84 | 2.99 |
|  | **Ramachandran Plot (%)** |  |  |  |  |  |
| Outliers | 0.00 | 0.00 | 0.00 | 0.00 | 0.00 | 0.00 |
| Allowed | 4.15 | 8.06 | 6.28 | 4.77 | 7.88 | 3.20 |
| Favored | 95.85 | 91.94 | 93.72 | 95.23 | 92.12 | 96.80 |
| Rotamer Outliers (%) | 0.00 | 0.00 | 0.00 | 0.00 | 0.00 | 0.00 |
|  | **Peptide Plane (%)** |  |  |  |  |  |
| Cis Proline/General | 0.0/0.0 | 0.0/0.0 | 0.0/0.0 | 0.0/0.0 | 0.0/0.0 | 0.0/0.0 |
| Twisted Proline/General | 0.0/0.0 | 0.0/0.0 | 0.0/0.0 | 0.0/0.0 | 0.0/0.0 | 0.0/0.0 |
| CaBLAM Outliers (%) | 3.96 | 3.14 | 2.89 | 3.45 | 7.84 | 5.26 |
|  | **Data** |  |  |  |  |  |
| Lengths (Å) | 63.07, 68.42, 94.07 | 90.38, 69.94, 82.85 | 87.81, 68.77, 102.63 | 71.48, 93.14, 106.13 | 102.22, 97.92, 66.71 | 67.35, 80.18, 106.90 |
| Angles (°) | 90.00, 90.00, 90.00 | 90.00, 90.00, 90.00 | 90.00, 90.00, 90.00 | 90.00, 90.00, 90.00 | 90.00, 90.00, 90.00 | 90.00, 90.00, 90.00 |
| Supplied Resolution (Å) | 3.8 | 4 | 3.7 | 4.3 | 4.4 | 4.5 |
| Resolution Estimates (Å) | **Masked** |  |  |  |  |  |
| d FSC (Half Maps; 0.143) | 3.8 | 3.9 | 3.7 | 4.0 | 4.1 | 4.1 |
| d 99 (Full/Half1/Half2) | 2.5/2.3/2.3 | 2.5/2.3/2.2 | 2.3/2.2/2.2 | 2.7/2.7/2.7 | 2.9/3.2/3.3 | 3.0/4.7/4.6 |
| d Model | 2.2 | 2.3 | 2.2 | 2.3 | 2.0 | 2.8 |
| d FSC Model (0/0.143/0.5) | 1.5/2.1/3.9 | 1.6/2.2/4.3 | 1.5/2.1/3.8 | 1.7/2.2/4.1 | 1.9/2.7/4.8 | 2.1/3.0/4.8 |
| Map Min/Max/Mean | 0.00/2.04/0.03 | 0.00/2.54/0.02 | 0.00/2.27/0.02 | 0.00/1.92/0.02 | 0.00/2.00/0.03 | 0.00/1.93/0.03 |
|  | **Model vs Data** |  |  |  |  |  |
| CC(Mask/Box/Peaks/Volume) | 0.72/0.68/0.60/0.73 | 0.61/0.64/0.57/0.62 | 0.72/0.74/0.70/0.73 | 0.74/0.65/0.52/0.74 | 0.65/0.62/0.46/0.65 | 0.64/0.56/0.37/0.64 |

