## Supplementary Materials for "Defining the Antigenic Topology and Prospective Binding Breadth of Vaccination-induced SARS-CoV-2 Neutralizing Antibodies"

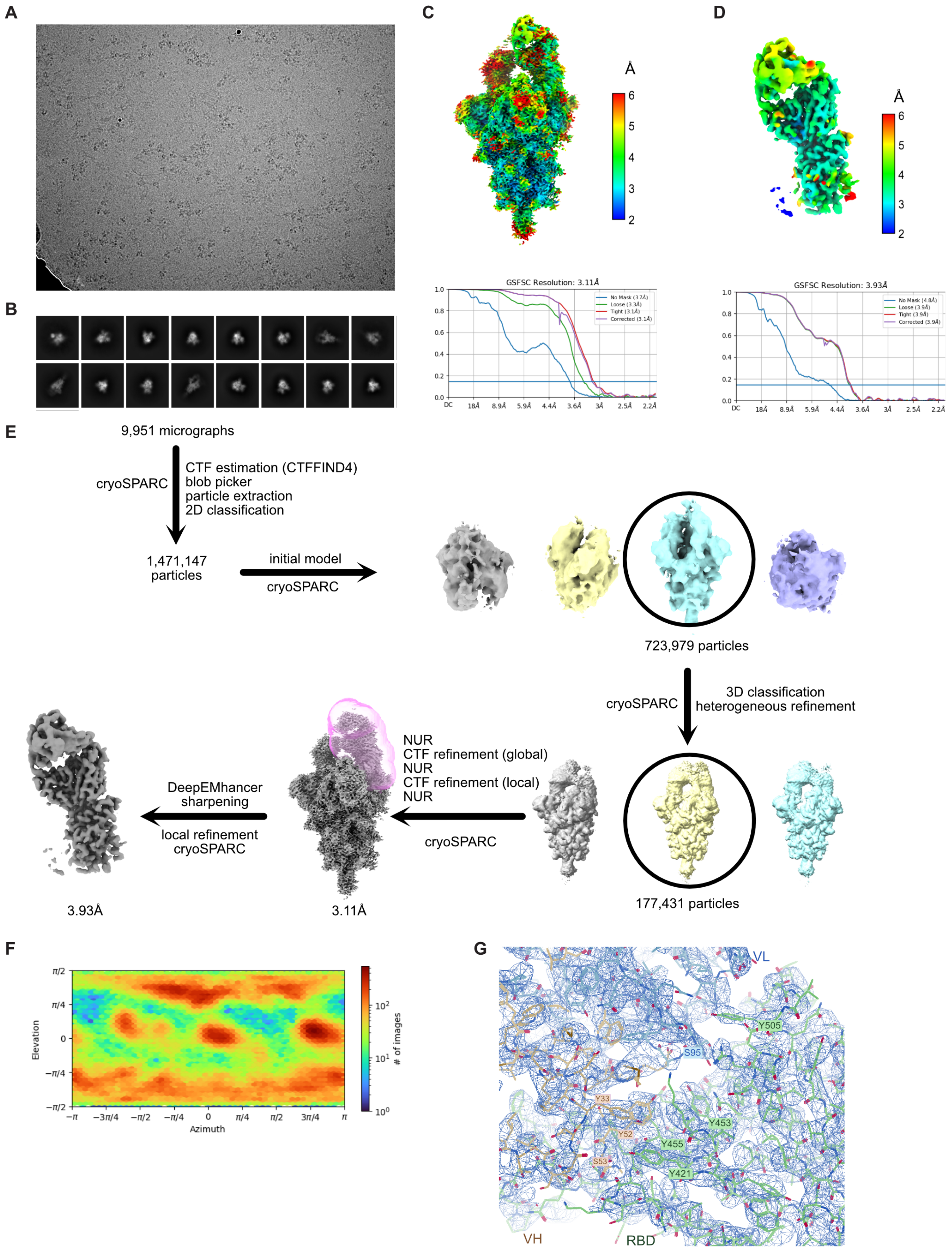


Figure 1—figure supplement 1.tiff

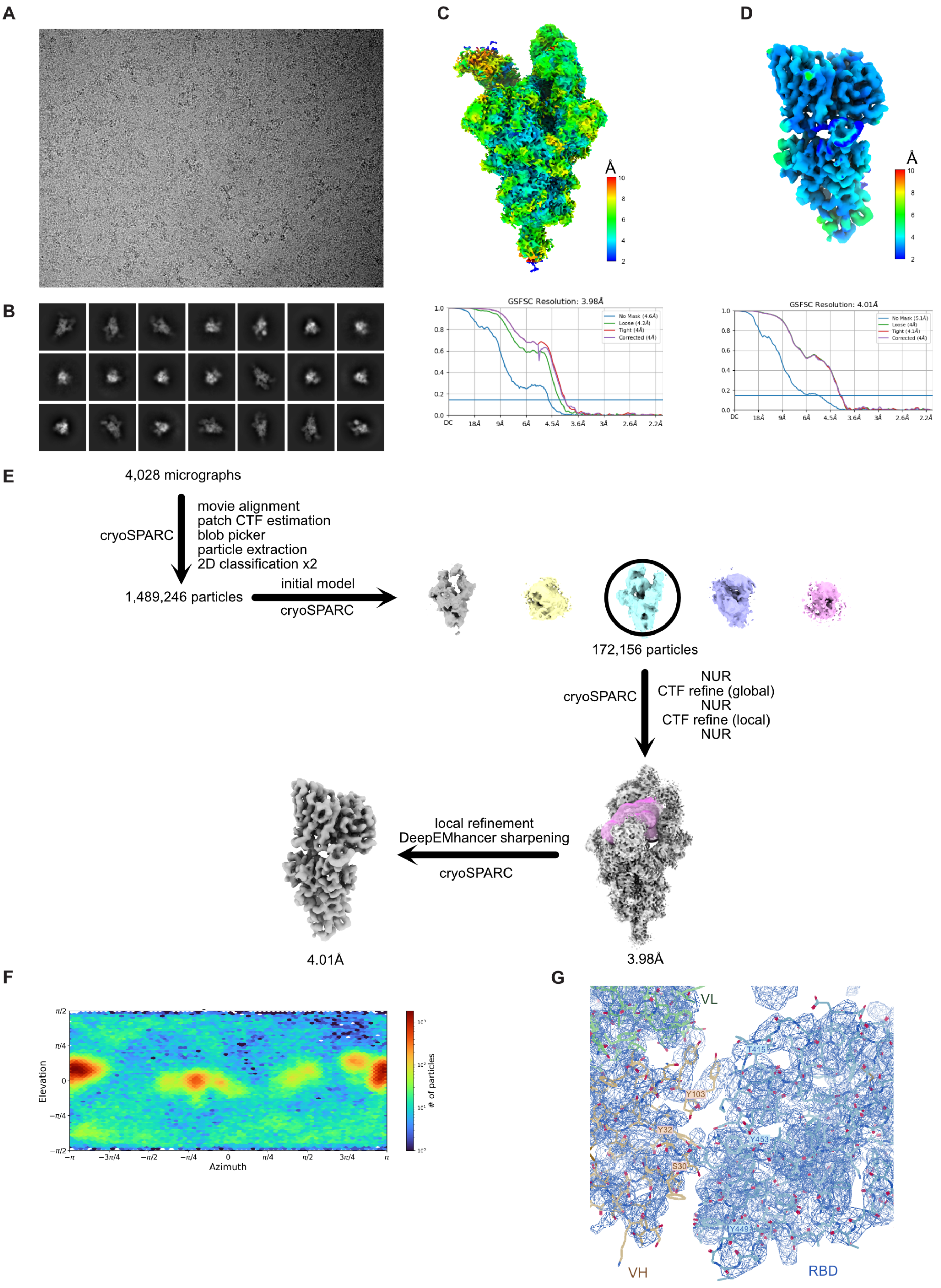


Figure 1—figure supplement 2.tiff

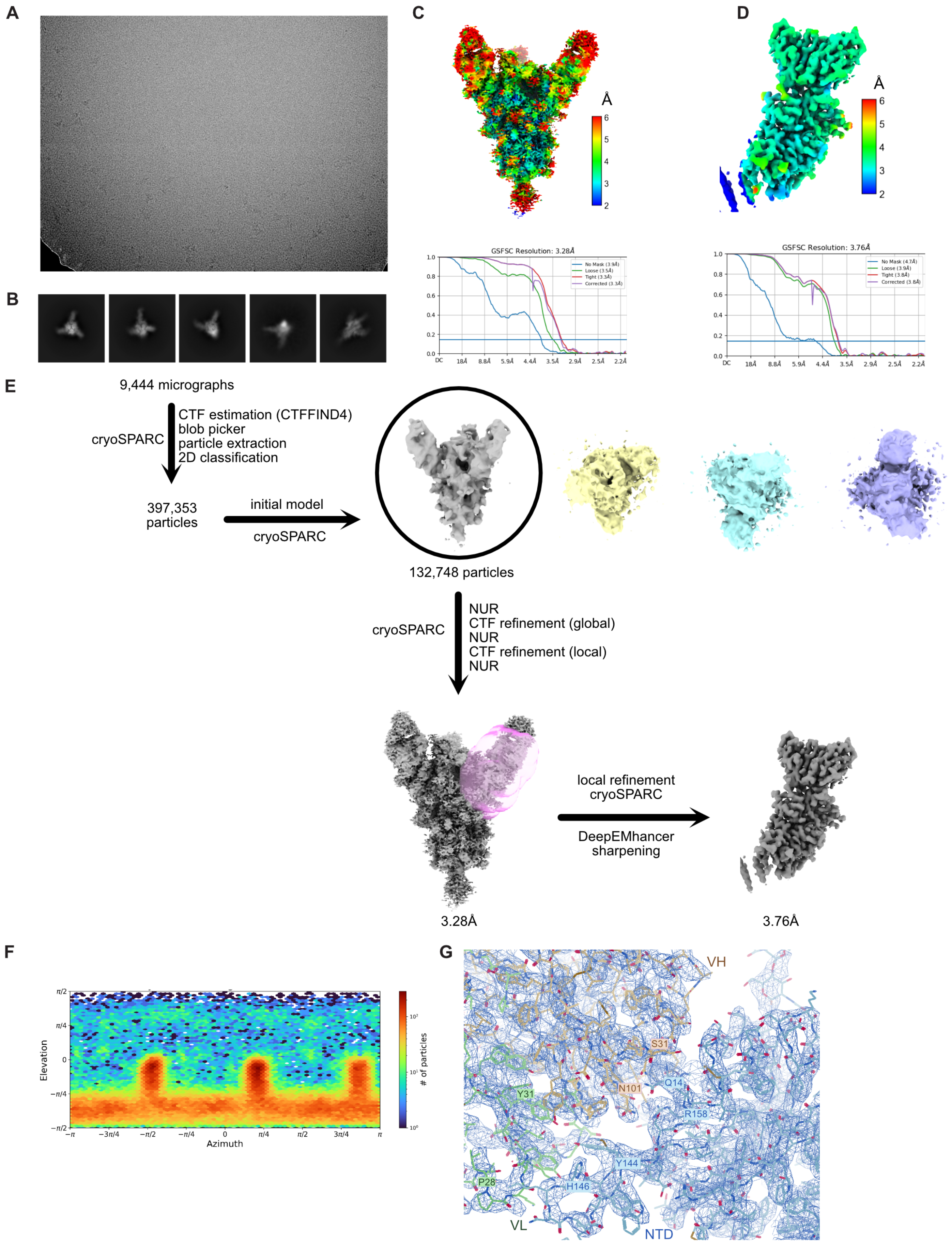


Figure 2—figure supplement 1.tiff

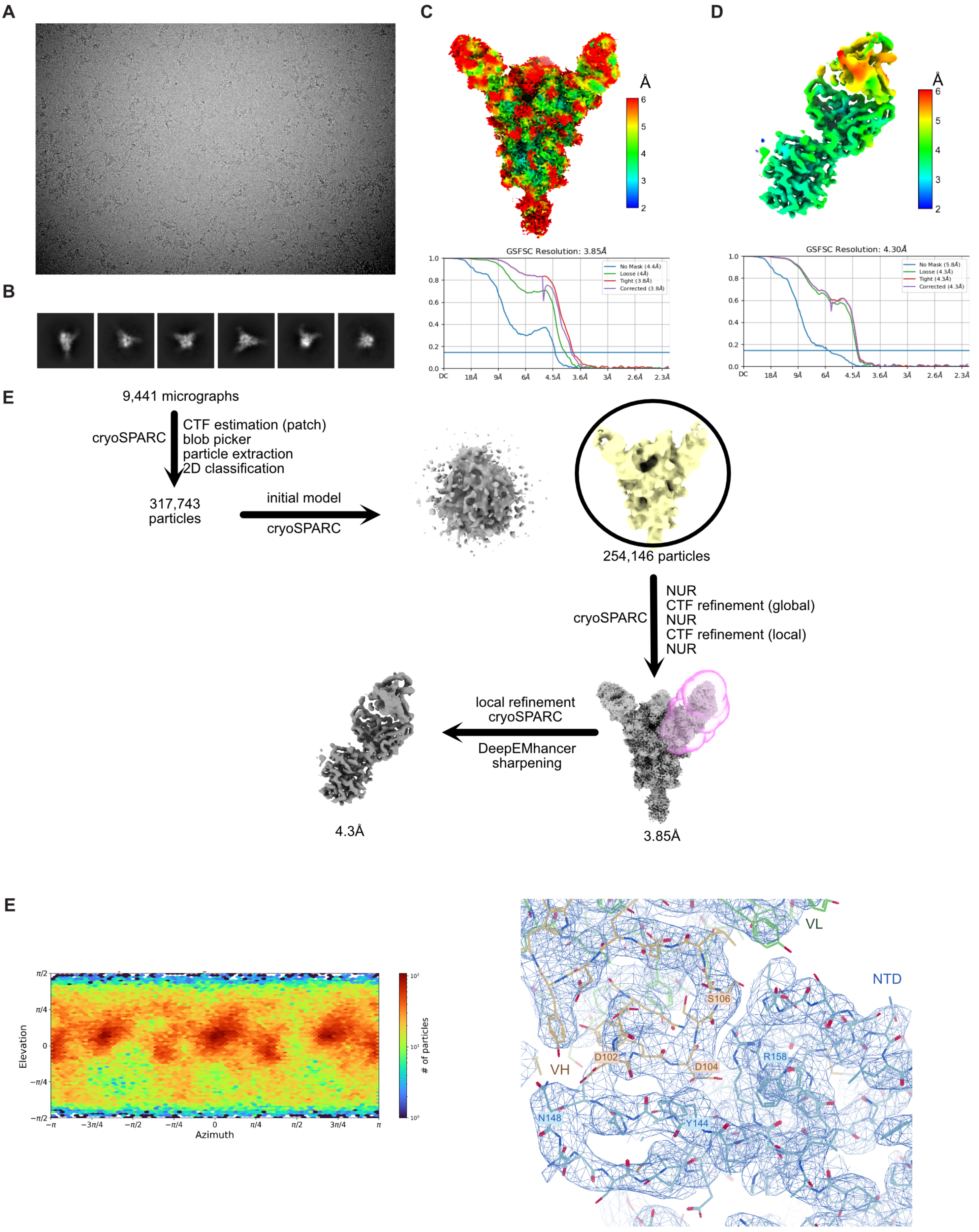


Figure 2—figure supplement 2.tiff

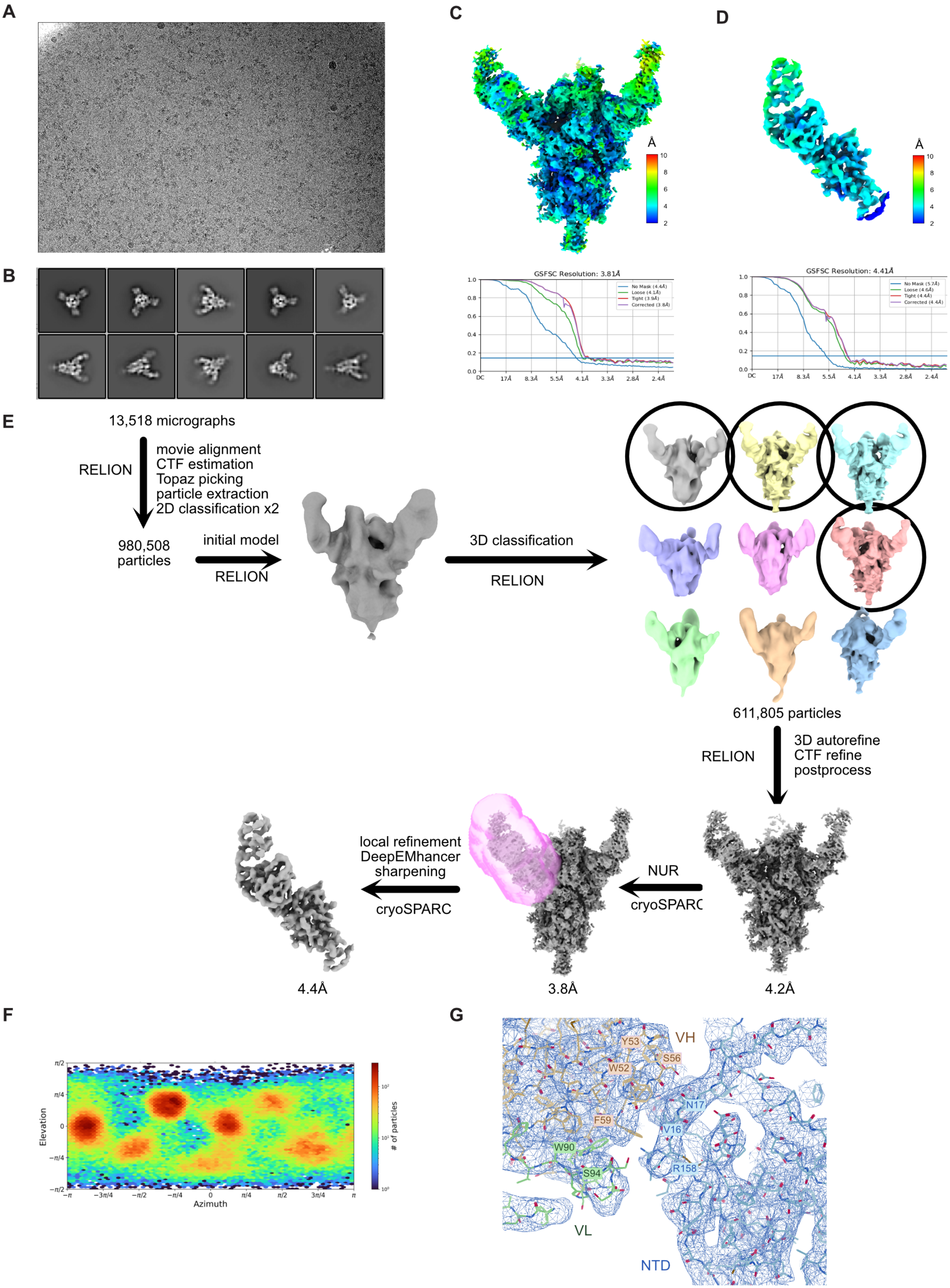


Figure 2—figure supplement 3.tiff

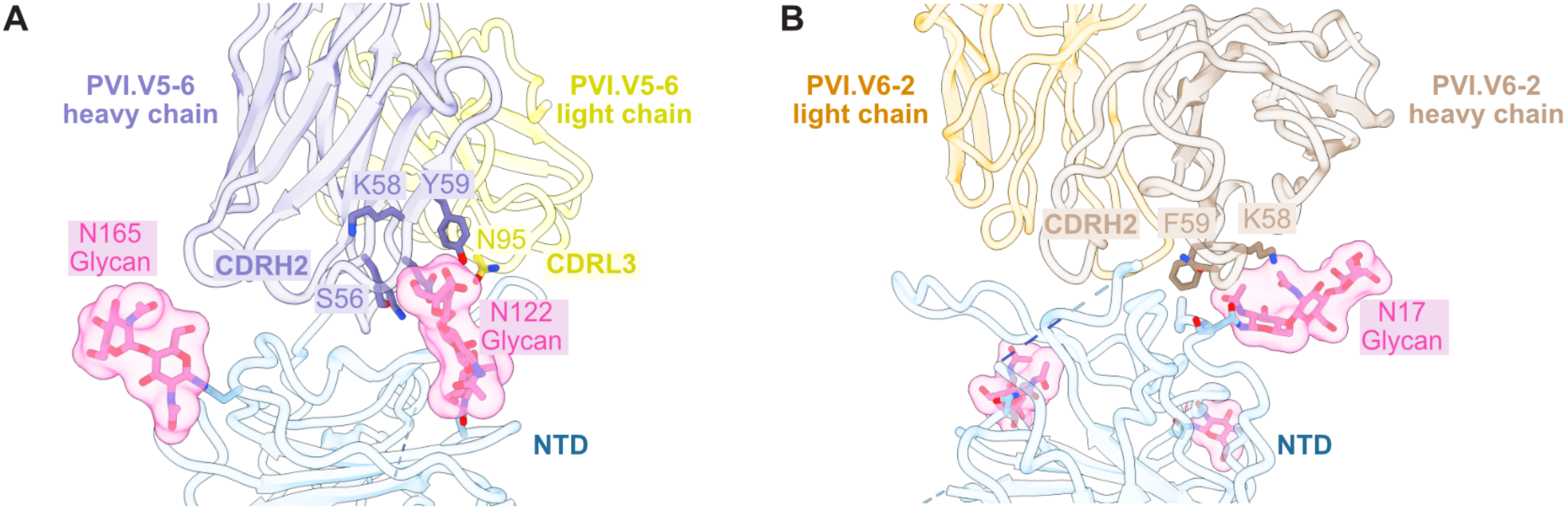


Figure 2—figure supplement 4.tiff

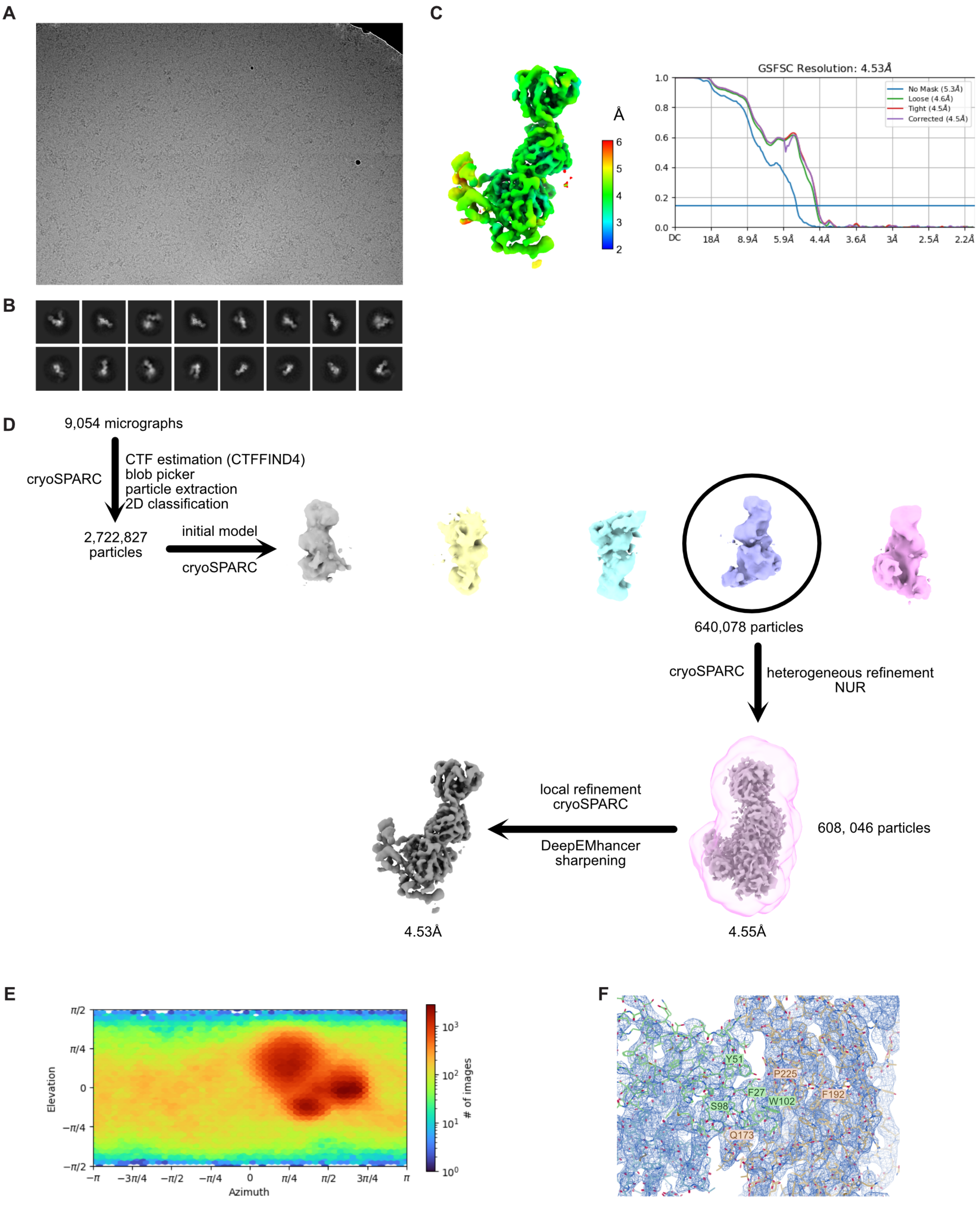


Figure 3—figure supplement 1.tiff

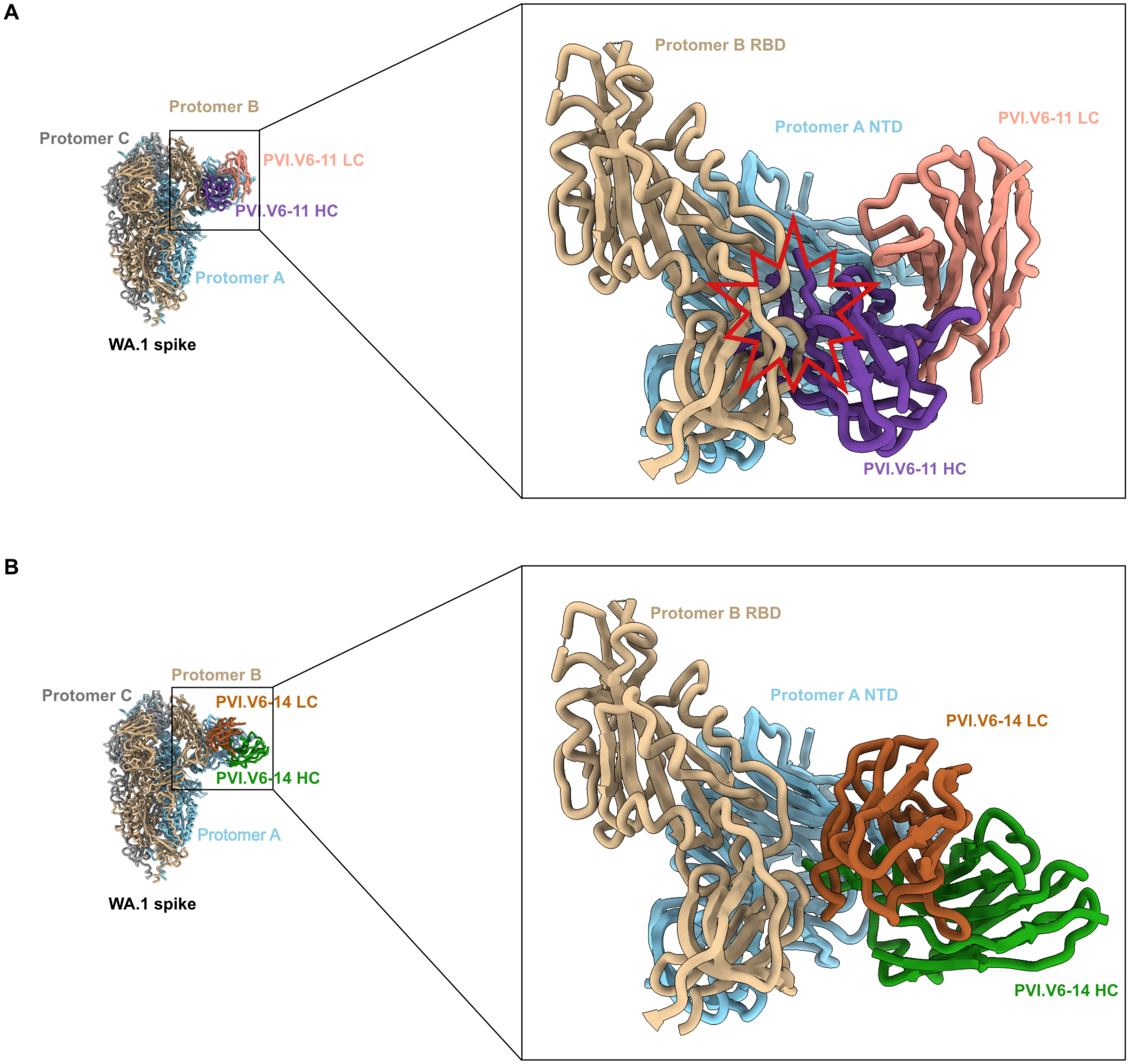


Figure 3—figure supplement 2.tiff

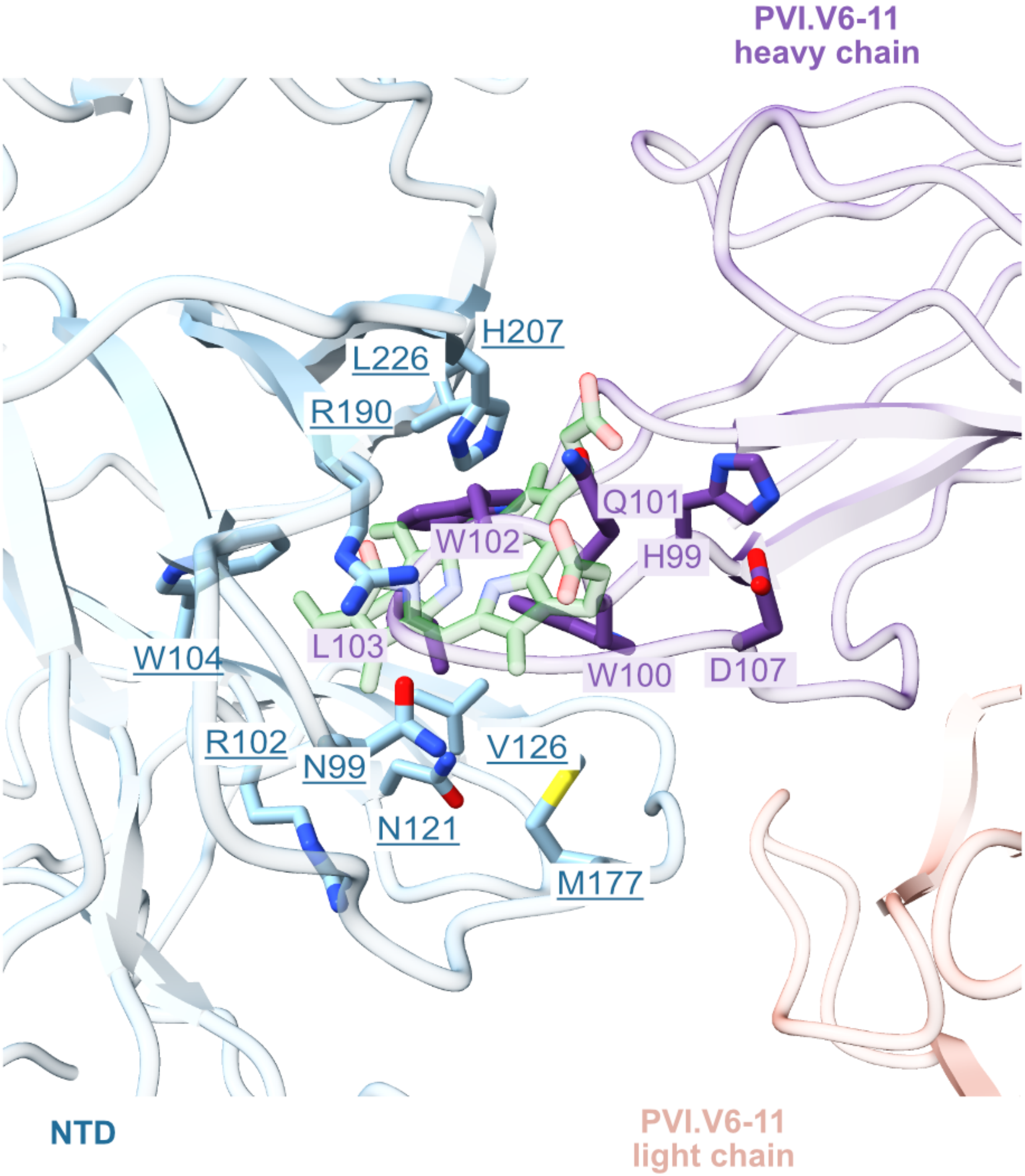


Figure 3—figure supplement 3.tiff

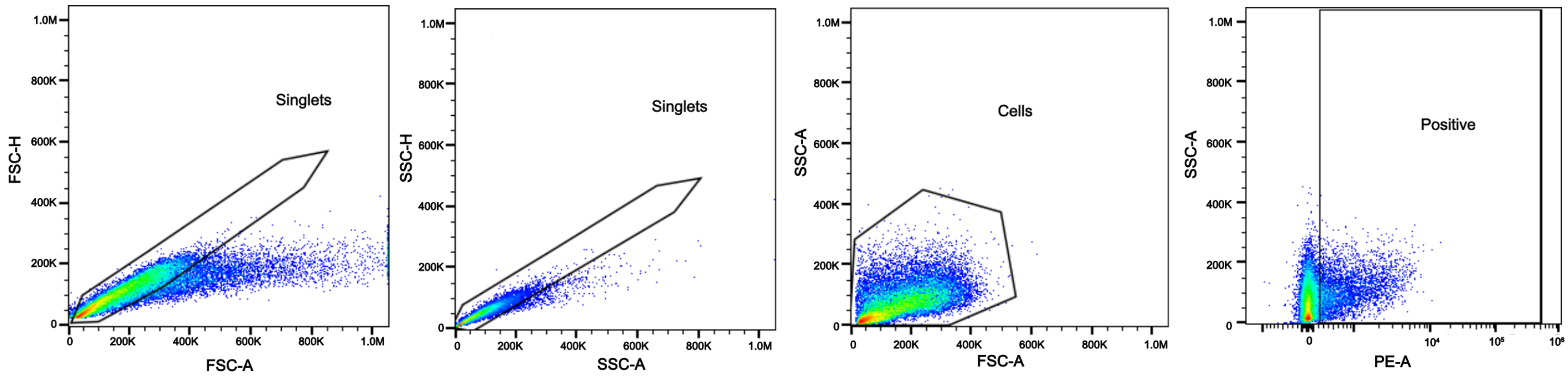


Figure 4—figure supplement 1.tiff

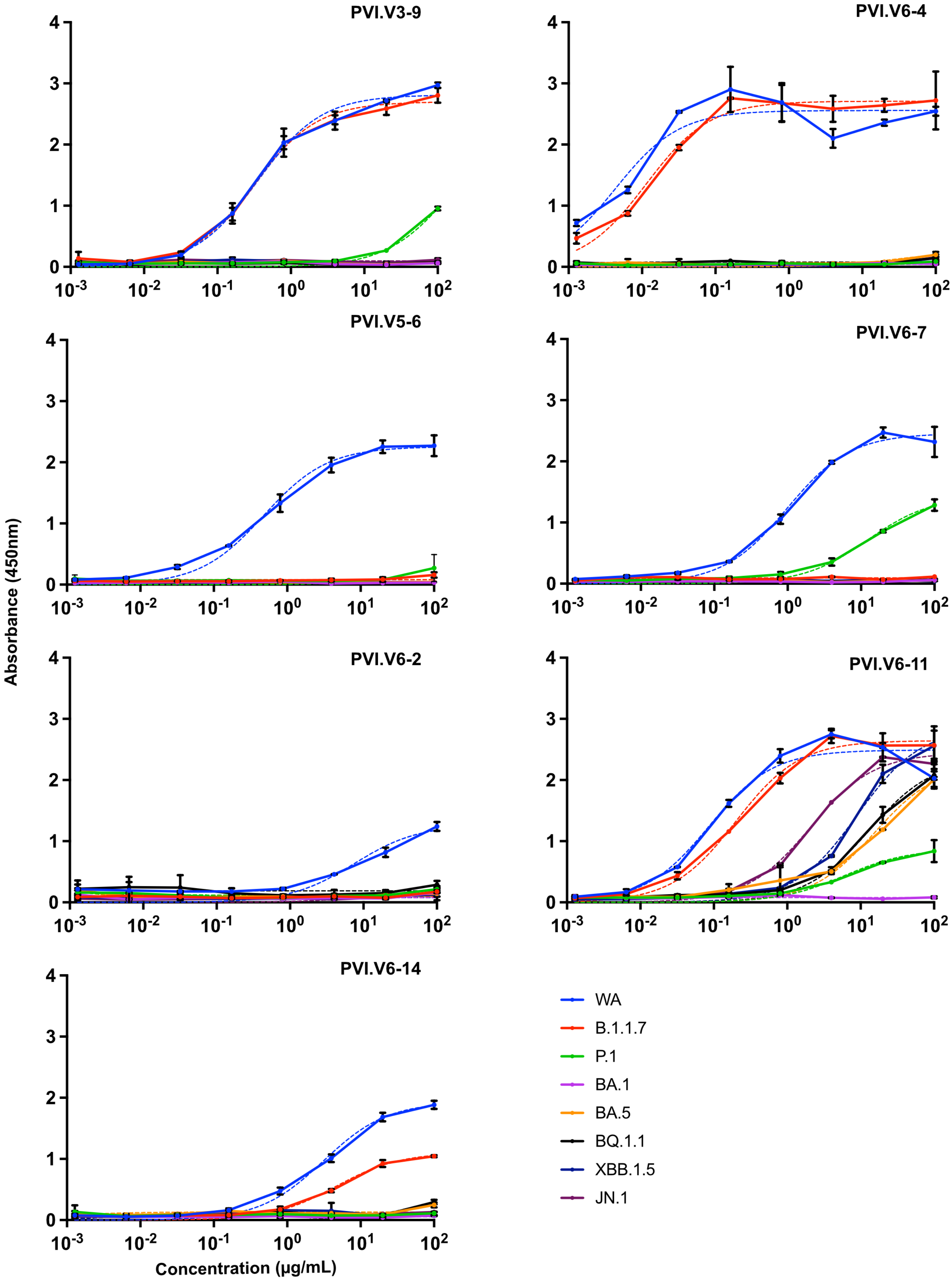


Figure 4—figure supplement 2.tiff

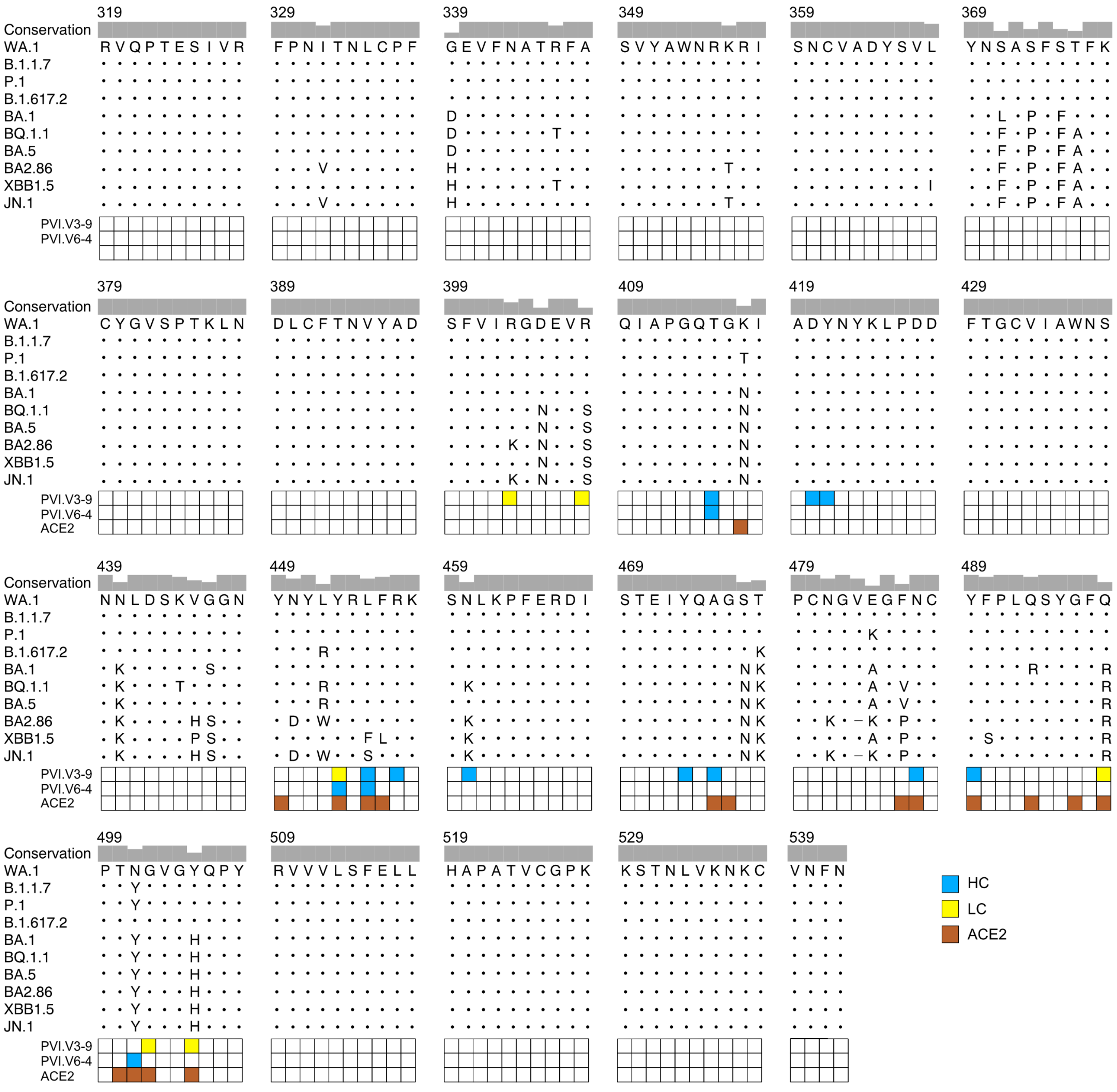

Figure 4—figure supplement 3.tiff

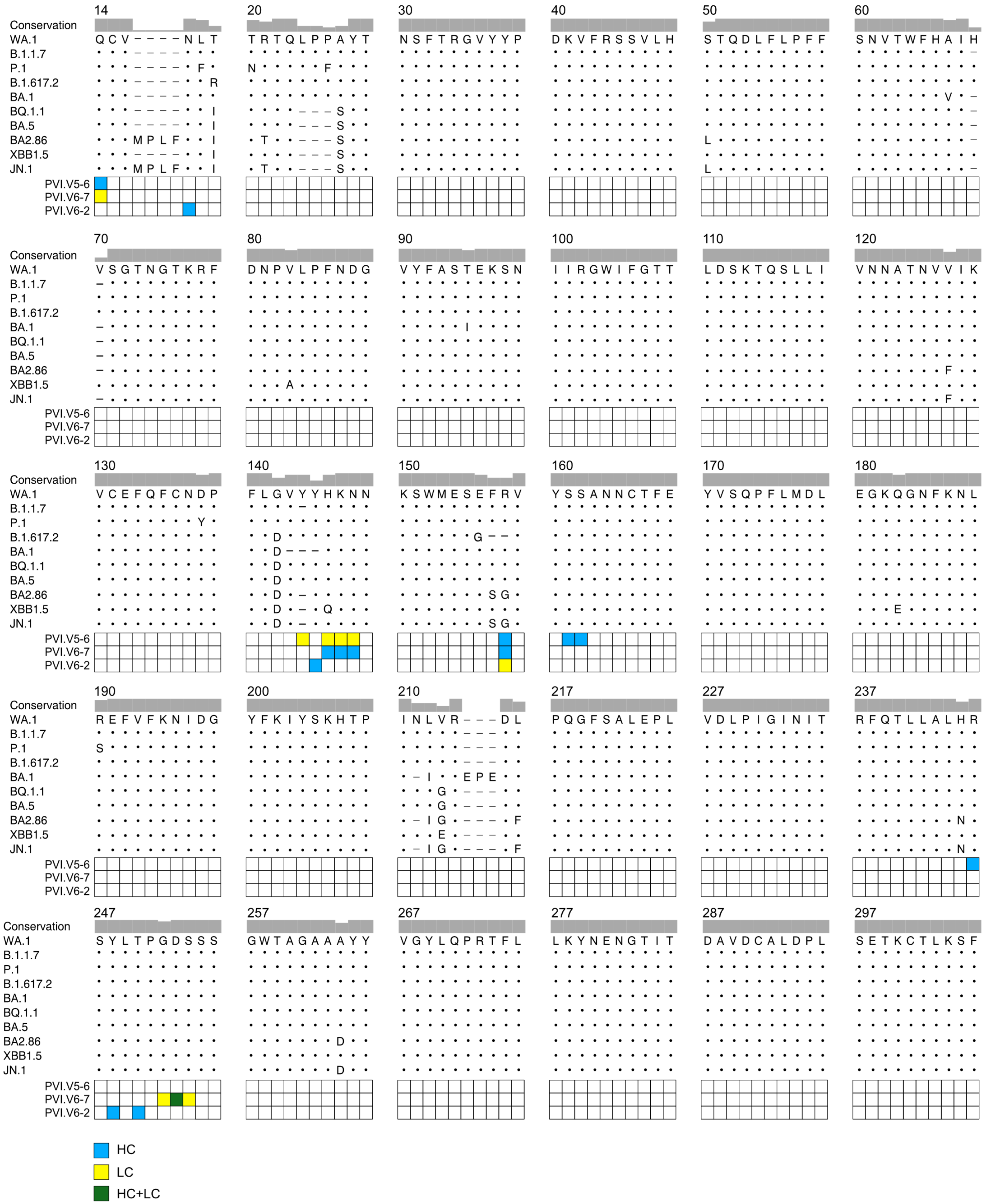


Figure 4—figure supplement 4.tiff

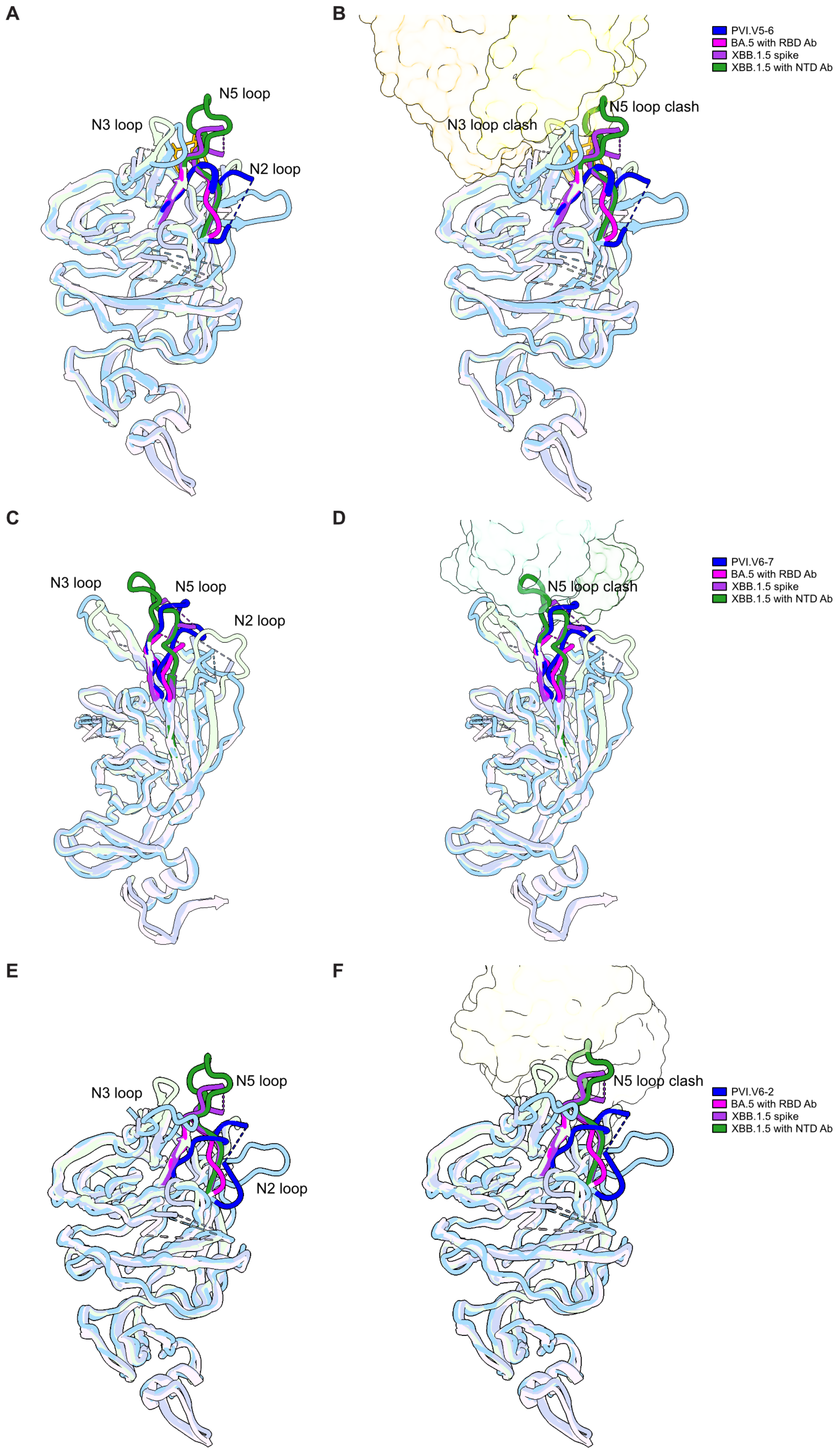


Figure 4—figure supplement 5.tiff

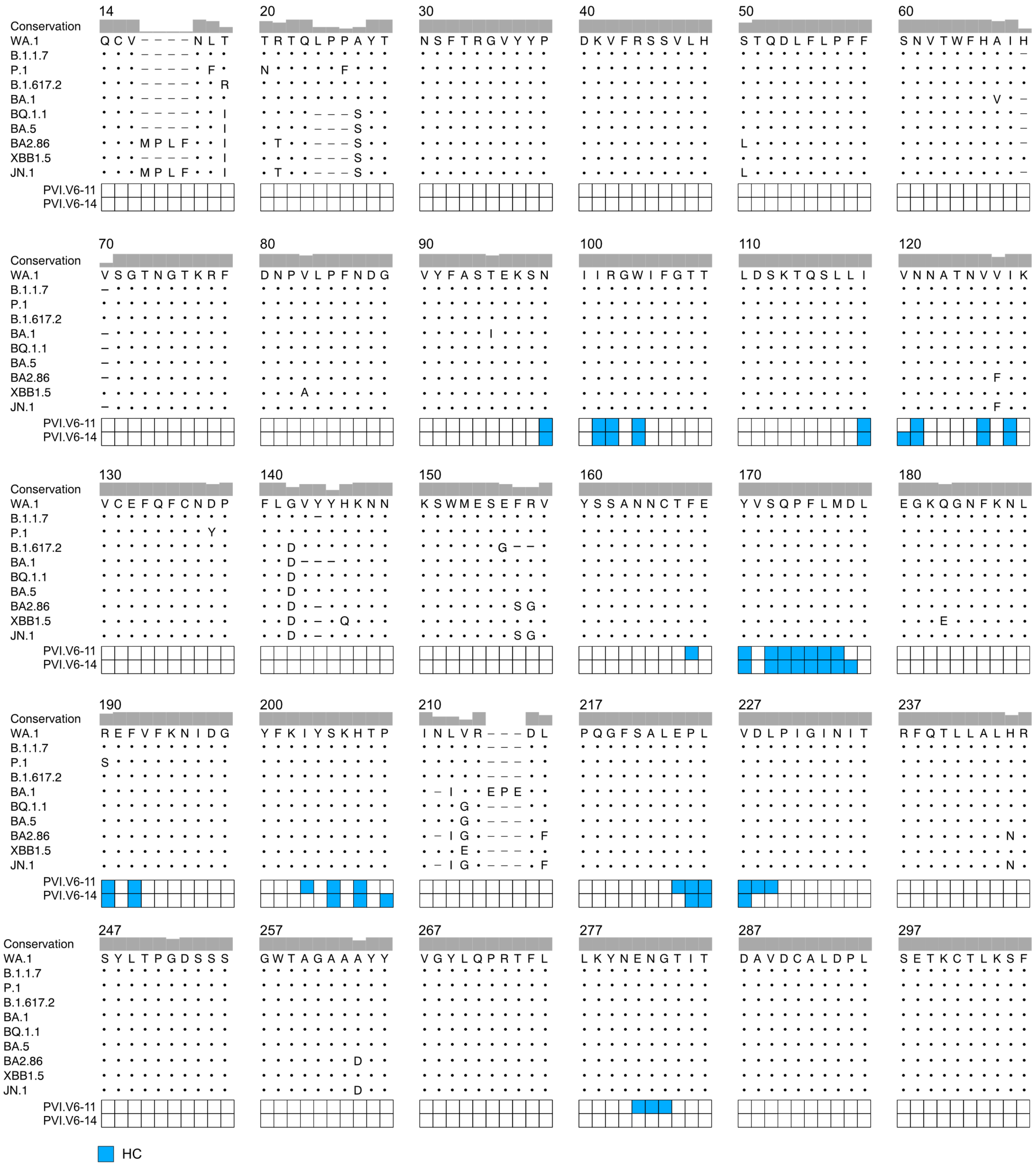

Figure 4—figure supplement 6.tiff
